# The recency and geographical origins of the bat viruses ancestral to SARS-CoV and SARS-CoV-2

**DOI:** 10.1101/2023.07.12.548617

**Authors:** Jonathan E. Pekar, Spyros Lytras, Mahan Ghafari, Andrew F. Magee, Edyth Parker, Jennifer L. Havens, Aris Katzourakis, Tetyana I. Vasylyeva, Marc A. Suchard, Alice C. Hughes, Joseph Hughes, David L. Robertson, Simon Dellicour, Michael Worobey, Joel O. Wertheim, Philippe Lemey

**Author notes:** These authors contributed equally (Jonathan E. Pekar, Spyros Lytras). These authors jointly supervised the work (David L. Robertson, Simon Dellicour, Michael Worobey, Joel O. Wertheim, Philippe Lemey).

## Abstract

The emergence of SARS-CoV in 2002 and SARS-CoV-2 in 2019 has led to increased sampling of related sarbecoviruses circulating primarily in horseshoe bats. These viruses undergo frequent recombination and exhibit spatial structuring across Asia. Employing recombination-aware phylogenetic inference on bat sarbecoviruses, we find that the closest-inferred bat virus ancestors of SARS-CoV and SARS-CoV-2 existed just ∼1–3 years prior to their emergence in humans. Phylogeographic analyses examining the movement of related sarbecoviruses demonstrate that they traveled at similar rates to their horseshoe bat hosts and have been circulating for thousands of years in Asia. The closest-inferred bat virus ancestor of SARS-CoV likely circulated in western China, and that of SARS-CoV-2 likely circulated in a region comprising southwest China and northern Laos, both a substantial distance from where they emerged. This distance and recency indicate that the direct ancestors of SARS-CoV and SARS-CoV-2 could not have reached their respective sites of emergence via the bat reservoir alone. Our recombination-aware dating and phylogeographic analyses reveal a more accurate inference of evolutionary history than performing only whole-genome or single gene analyses. These results can guide future sampling efforts and demonstrate that viral genomic fragments extremely closely related to SARS-CoV and SARS-CoV-2 were circulating in horseshoe bats, confirming their importance as the reservoir species for SARS viruses.

## Introduction

Horseshoe bats (*Rhinolophus spp.*) are the main hosts of the *Sarbecovirus* subgenus^1^ (genus *Betacoronavirus,* family *Coronaviridae*) and, likely via virus transmission through a transient intermediate species, were the source of SARS-CoV (referred to hereafter as SARS-CoV-1, now extinct) and SARS-CoV-2^2–4^. Since the emergence of SARS-CoV-1 in Guangdong Province in 2002 and SARS-CoV-2 (jointly referred to as the SARS-CoVs) in Hubei Province in 2019, there has been increased sampling of sarbecoviruses in horseshoe bats, which has contributed to our understanding of sarbecovirus diversity and their geographical spread.

Genome-wide sequence identity is typically used to compare bat sarbecoviruses to the SARS-CoVs, but because coronaviruses frequently recombine^5,6^, whole genome identity of these bat viruses and the SARS-CoVs does not adequately reflect their complex evolutionary histories. For example, the most recent common ancestor (MRCA) of SARS-CoV-2 and a closely-related bat virus genome sequence might be estimated at ∼20 years before 2019. However, when analyzing subgenomic, non-recombinant regions (NRRs) we will inevitably find some NRRs with older and some with younger MRCAs than this whole-genome estimate, which is effectively a weighted average of NRR tMRCAs. Because recombination is so rampant among bat sarbecoviruses, we likely would have had to have sampled the sarbecovirus from the individual bat involved in each cross-species transmission event leading to the SARS-CoVs (which we refer to as the “direct ancestor”; Extended Data Fig. 1), to have sampled a bat virus that had no recombination in its genome relative to these SARS-CoVs. What we have, instead, is a relatively small number of bat sarbecovirus genome sequences, gathered across different times and places. Hence, it is necessary to identify the NRRs of sarbecoviruses—genomic regions within which there is little to no detectable signal of recombination among the analyzed genomes and representing, to the extent possible, a single evolutionary history. These regions provide the clearest insights into the evolutionary history of SARS-CoV-1 and SARS-CoV-2 and the origins of their respective outbreaks.

Sarbecoviruses exhibit substantial geographic structuring linked to the ranges of their host horseshoe bat species^2–45,66^, but where and when their ancestors circulated is poorly understood. The sarbecoviruses have an ancient origin^7–9^, whose timing is difficult to infer due to substitution saturation, the repeated occurrence of substitutions at the same site resulting in biased estimation of older divergence in viral phylogenies^10,11^. This feature of viruses complicates dating estimates of deep nodes in the sarbecovirus phylogenies and could falsely suggest that their ancestors circulated more recently than they actually did^12^.

Here, we separately analyze the recombination patterns and respective evolutionary histories of SARS-CoV-1 and its closely related viruses (referred to as SARS-CoV-1-like viruses) and SARS-CoV-2 and its closely related viruses (referred to as SARS-CoV-2-like viruses). We estimate the separate sets of NRRs for the SARS-CoV-1-like and SARS-CoV-2-like viruses. For each NRR we utilize a molecular clock rate specific for that portion of the genome and determine the MRCA of the SARS-CoV and the most closely related bat virus, or set of viruses, among currently sampled sarbecoviruses for the NRR. We refer to this MRCA as the “closest-inferred ancestor” for that NRR because it is the parent node of the SARS-CoV in the phylogeny (Extended Data Fig. 1) and is limited by the current sample set of bat sarbecoviruses.

We show that the available sequence data provide evidence of the closest-inferred bat virus ancestors of both SARS-CoV-1 and SARS-CoV-2 circulating in bats in the years immediately preceding the emergence of each SARS-CoV. By performing a phylogeographic analysis of these sarbecoviruses and accounting for substitution saturation^7,8^, we demonstrate that the sarbecoviruses spread at a rate similar to their bat hosts and that, based on current sampling, both SARS-CoV-1 and SARS-CoV-2 likely emerged far from their direct bat virus ancestors.

### Inference of non-recombinant regions

To identify the recombination patterns of the sarbecoviruses, we aligned the available 167 full-genome sarbecovirus sequences (Supplementary Data File 1), including SARS-CoV-1 and SARS-CoV-2. We used GARD^13^ to independently identify recombination breakpoints on genomes closely related to SARS-CoV-1 and SARS-CoV-2 (Extended Data Fig. 2). These breakpoints identify 31 putative NRRs for the SARS-CoV-1-like viruses, with a median length of 866 nucleotides (nt) and a range of 309 to 2195 nt, and 27 putative NRRs for the SARS-CoV-2-like viruses, with a median length of 970 nt and a range of 252 to 2836 nt (Fig. 1a).

**Figure 1.**
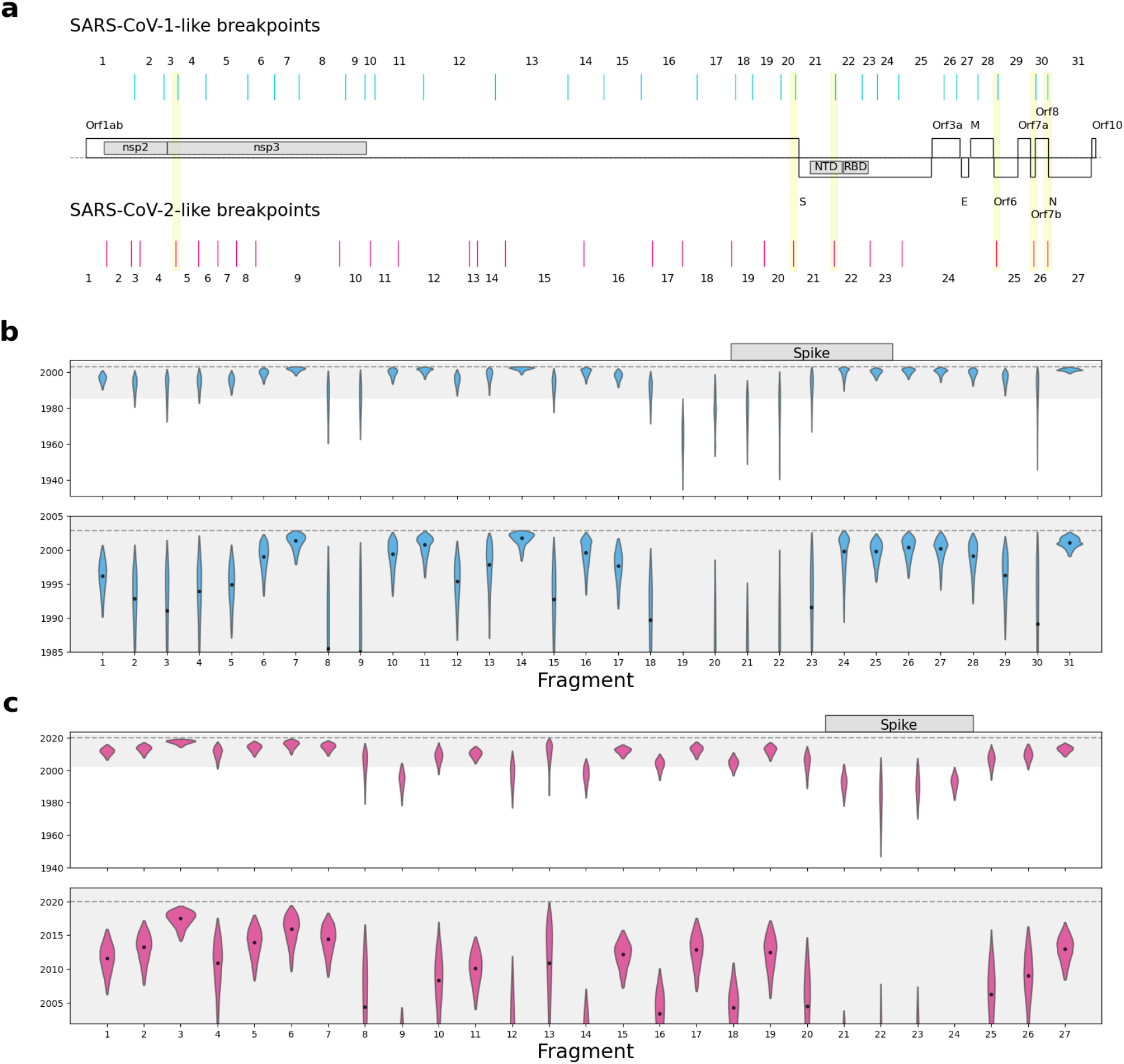
Recombination patterns and dating of the closest-inferred bat virus ancestor of SARS-CoV-1 and SARS-CoV-2 for each NRR. **a,** Distribution of SARS-CoV-1-like and SARS-CoV-2-like breakpoints across the sarbecovirus genome alignment. Schematic of major open-reading frames is displayed in the middle of the figure. Gene domains of interest are also shown within the schematic. Breakpoint pairs with significantly short distances (<10% point of distribution) are highlighted in yellow. Numbers indicate NRRs resulting from breakpoints. The alignment is 31,243 nt long. **b,** Time of the closest-inferred bat virus ancestor of SARS-CoV-1 across the 31 NRRs of the SARS-CoV-1-like viruses. The 20 years before the emergence of SARS-CoV-1 is shown in gray in the upper panel and zoomed-in upon in the lower panel. c, Time of the closest-inferred bat virus ancestor of SARS-CoV-2 across the 27 NRRs of the SARS-CoV-2-like viruses The 20 years before the emergence of SARS-CoV-2 is shown in gray in the upper panel and zoomed-in upon in the lower panel. The dashed line in each panel is the date of the earliest sampled human case of each SARS-CoV (16 November 2002 for SARS-CoV-1; 24 December 2019 for SARS-CoV-2). The violin plots correspond to the 95% highest posterior density (HPD) intervals and the dot within each violin plot indicates the median. NRRs corresponding to the Spike gene are indicated in the top panels of **b-c**.

To examine whether the inferred positions of recombination breakpoints of the SARS-CoV-1-like and SARS-CoV-2-like NRR analyses were closer to each other than expected, we simulated randomly distributed recombination breakpoint sets along the alignment for the SARS-CoV-1-like and SARS-CoV-2-like viruses (30 and 26 breakpoints, respectively) and compared the distribution of distances between the two breakpoint sets. The empirical breakpoint sets had a mean genomic distance of 307 nt (90% CIs: 204-463 nt) separating the SARS-CoV-1 and SARS-CoV-2 breakpoints, whereas the simulated distances had a mean of 540 nt (90% CIs: 334-872 nt) (Extended Data Fig. 3). Additionally, there were six breakpoint pairs between the two virus sets that were in the smallest 10% of the observed distance distribution (15 to 78 nt apart; Fig. 1a), indicating that hotspots of recombination are likely shared between the two sarbecovirus clades (*e.g.*, 5’ end of nsp3, upstream of Spike, and other accessory genes).

### Sarbecovirus genomes provide evidence for a very recent bat ancestor of SARS-CoV-1 and SARS-CoV-2

The evolutionary history of sarbecoviruses can only be accurately inferred when focusing on NRRs rather than entire genomes or specific genes because these viruses, as well as their ancestors, have undergone extensive recombination^4,6,14^. Here, we examine when the closest-inferred bat viruses ancestral to SARS-CoV-1 and SARS-CoV-2 circulated. These ancestors are represented by the MRCA of each SARS-CoV and its most closely related bat virus (or viruses), which is akin to the parent node of each SARS-CoV in its respective phylogeny.

There is insufficient temporal signal when calibrating a molecular clock using tip dating with sarbecoviruses sampled from bats and pangolins (*Manis javanica*)^6^, likely as a consequence of limited sampling across space and time. Therefore, we used SARS-CoV-1 genomes to identify a suitable rate prior for the molecular clock in our time-scaled phylogenetic analyses (Extended Data Fig. 1; see Methods). The inferred substitution rates served as NRR-specific rate priors for subsequent Bayesian phylogenetic inference of SARS-CoV-1 and the 139 SARS-CoV-1-like viruses (Extended Data Fig. 4, Supplementary Table 1). The median time of the closest-inferred bat virus ancestor of SARS-CoV-1 within each NRR varied by several decades (Fig. 1b), and the median of these medians is 1996. However, the most recent time of the closest-inferred ancestor across all NRRs of SARS-CoV-1 was 2001 (95% highest posterior density [HPD]: 1998–2002; NRR 14 [Orf1b]; 1065 nt), only one year before the emergence of SARS-CoV-1 in 2002. The viruses most similar to SARS-CoV-1 within this NRR were YNLF_31C and YNLF_34C, sampled in Yunnan, China in 2013. Molecular clock inference informed by narrower rate priors resulted in times of the closest-inferred ancestors for several NRRs closer to the emergence of SARS-CoV-1 (Extended Data Fig. 5a, Supplementary Table 3). Our findings were consistent when we included a more divergent human SARS-CoV-1 genome along with a masked palm civet (*Paguma larvata*) SARS-CoV-1 genome (Extended Data Fig. 5a, Supplementary Table 3).

We similarly used the evolutionary rate of SARS-CoV-2 across the inferred NRRs of the SARS-CoV-2-like viruses (Extended Data Fig. 4, Supplementary Table 2) as priors for the phylogenetic analysis of the NRRs of the SARS-CoV-2-like viruses, comprising SARS-CoV-2 and the 26 SARS-CoV-2-like genomes. The time of the closest-inferred bat virus ancestor of SARS-CoV-2 ranged from several years to several decades preceding late 2019 — when SARS-CoV-2 was introduced into humans — across the 27 NRRs (Fig. 1c). Although the median time of the closest-inferred ancestor of SARS-CoV-2 was in 2009, the most recent time of the closest-inferred ancestor across all NRRs was in 2017 (95% HPD: 2014–2019; NRR 3 [Orf1a]; 259 nt), only two years prior to its introduction into humans^15^. The genome most similar to SARS-CoV-2 within this NRR is RmYN02, sampled in Yunnan, China in 2019. We note that although NRR 3 is only 259 nt, NRR 6 is 573 nt and has a similar time of the closest-inferred ancestor of 2015 (95% HPD: 2009–2019). Within NRR 6, the genome most related to SARS-CoV-2 is BANAL-20-103, sampled in Laos in 2020. A narrower clock rate prior results in similar inferred dates (Extended Data Fig. 5b, Supplementary Table 4).

The times of closest-inferred ancestor of SARS-CoV-1 across the NRRs are nearer to the date of human emergence than those for SARS-CoV-2 (Fig. 1b,c): 14 NRRs of the SARS-CoV-1-like viruses have a median time of the closest-inferred ancestor of SARS-CoV-1 within 5 years of the emergence of SARS-CoV-1, whereas only 3 NRRs of the SARS-CoV-2-like viruses have a closest-inferred ancestor of SARS-CoV-2 within 5 years of the emergence of SARS-CoV-2. Both sets of viruses consistently have more divergent fragments across the Spike gene (Fig. 1b). The oldest time of the closest-inferred bat virus ancestor within a single NRR of SARS-CoV-1 and SARS-CoV-2 was, respectively, 1963 (95% HPD: 1934–1985; NRR 19 [Orf1b]) and 1982 (95% HPD: 1946–2007; NRR 22 [Spike]), with each approximately 40 years prior to the emergence of their corresponding SARS-CoV. Notably, only NRR 19 among the SARS-CoV-1-like viruses has a median time of the closest-inferred ancestor more than 50 years earlier than the most recently sampled SARS-CoV-1-like virus.

Accurate estimation of the ages of deeper nodes in the SARS-CoV phylogenies require complex models accommodating high levels of substitution saturation^7,11^. We restricted our analyses to the most recent nodes of the tree. Our inferred ages for the closest-inferred ancestors of each of the SARS-CoVs across the NRRs indicate that several of the published sarbecovirus genomes include non-recombinant fragments that are descendant from viruses that circulated likely only a few years before the emergence of SARS-CoV-1 and SARS-CoV-2.

### The recombinant common ancestors became more similar to SARS-CoV-1 and SARS-CoV-2 with increased sampling

We quantified the extent to which the publication of more viruses closely related to SARS-CoV-1 and SARS-CoV-2 has contributed toward our understanding of the ancestor of each of these human viruses. The respective recombinant common ancestor (“recCA”) of SARS-CoV-1 and SARS-CoV-2 can be understood as the aggregate of the closest-inferred ancestors of each of the SARS-CoVs across each of its NRRs^15^, with each closest-inferred ancestor existing at different points in time and with potentially different viruses most closely related to the SARS-CoV. The recCA therefore accounts for the closest relative(s) across all non-recombinant segments. We inferred the genome of the recCA of each SARS-CoV over time, progressively adding published genomes to the reconstruction based on their publication date. Then, we calculated the similarity of each recCA to its respective SARS-CoV as a function of the publication date to examine how additional closely related genomes can affect ancestral reconstruction and reflect our increasing understanding of sarbecoviruses over time (Fig. 2a).

**Figure 2.**
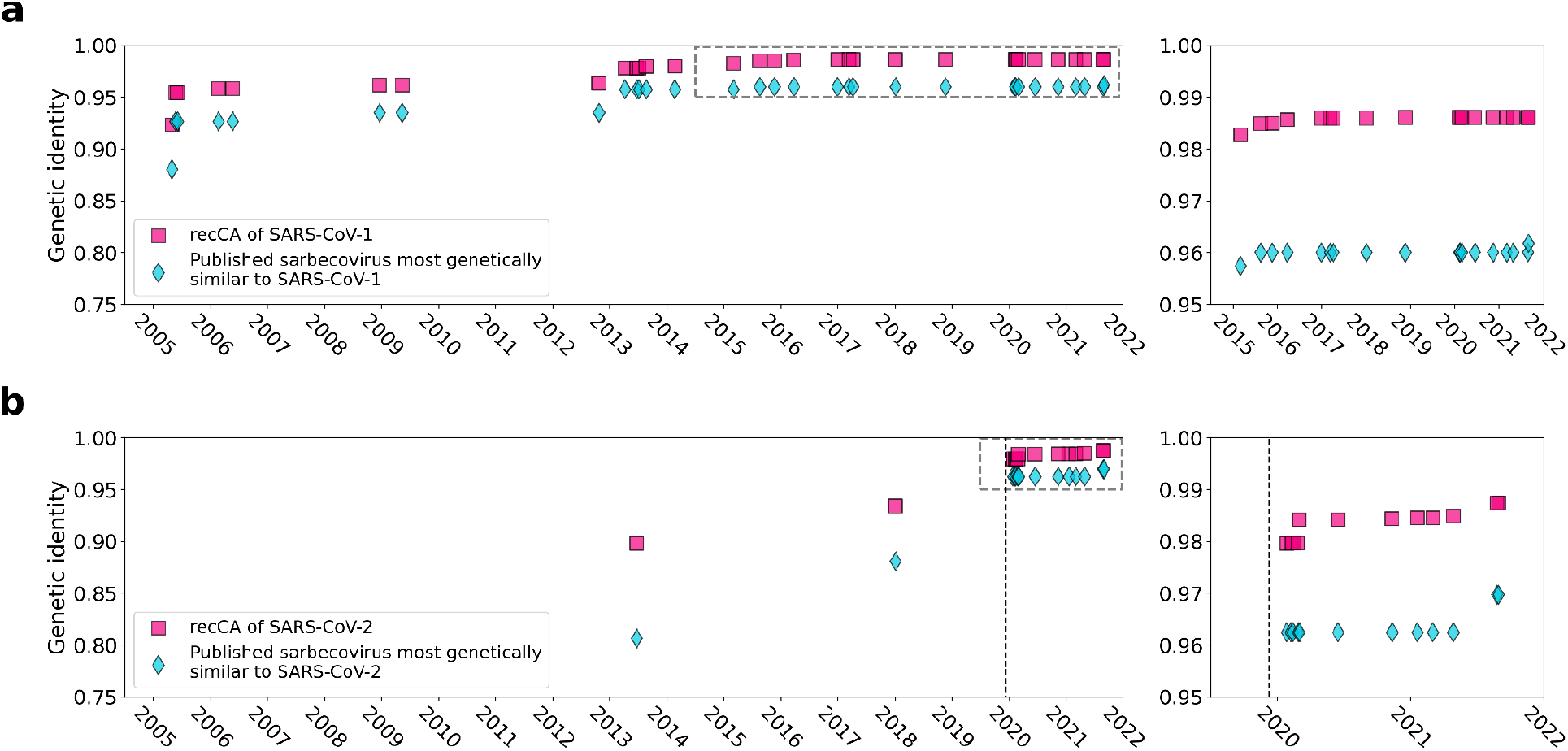
The recCA (recombinant common ancestor) and the most closely related sarbecovirus over time. **a,** The similarity of the recCA of SARS-CoV-1 and the most closely related sarbecovirus in the SARS-CoV-1 viruses to SARS-CoV-1 as a function of time. **b,** The similarity of the recCA of SARS-CoV-2 and the most closely related sarbecovirus in the SARS-CoV-2 viruses to SARS-CoV-2 as a function of time. The right panels for **a** and **b** are zoomed-in panels of the dashed box in the left panels. The dashed vertical line in **b** is the sampling date of the earliest sampled genome of SARS-CoV-2.

We observe that, as more genomes were published, the SARS-CoV-1 recCA and SARS-CoV-2 recCA became more similar to SARS-CoV-1 and SARS-CoV-2, respectively (Fig. 2). The full bat virus genomes most similar to SARS-CoV-1 and SARS-CoV-2 are, respectively, YN2020E (96.2% genetic identity to SARS-CoV-1) and BANAL-20-52 (96.8% genetic identity to SARS-CoV-2), both published in 2021. The genetic identity of the recCA of SARS-CoV-1 with SARS-CoV-1 exceeded the genetic identity of YN2020E with SARS-CoV-1 by 2013. The genetic identity of the recCA of SARS-CoV-2 with SARS-CoV-2 exceeded the genetic identity of BANAL-20-52 with SARS-CoV-1 in 2020, coinciding with the publication of the full genome of RaTG13 (96.2% genetic identity to SARS-CoV-2) in January 2020. The recCA of each SARS-CoV therefore exceeded the genetic similarity of the most closely related full genome before it was actually sampled.

Furthermore, the recCAs continued to become more similar to their corresponding SARS-CoV in the subsequent years. For example, the recCA of SARS-CoV-2 continued to increase in similarity after RaTG13 and before BANAL-20-52 were published, despite the interim published genomes sharing less overall genetic identity with SARS-CoV-2 than RaTG13 (Fig. 2b). This increase in similarity is the result of these genomes containing fragments that descend from a more closely related ancestor despite the rest of the genome being more divergent from SARS-CoV-2 than RaTG13. These results can therefore be observed at the level of individual NRRs: sequentially published genomes increase the genetic identity of some of the NRRs of the recCA to the matching regions of SARS-CoV-2 (Extended Data Fig. 6). However, not all NRRs of the recCA exhibit increases in genetic identity to SARS-CoV-2 with additionally published genomes, and occasionally there is no increase of genetic identity to SARS-CoV-2 across any of the NRRs of the recCA when a new genome is published. Regardless, our results show that, by the end of 2022, the inferred recCA of each SARS-CoV is very similar to these human viruses, with the recCA of SARS-CoV-2 sharing 98.8% genetic identity with SARS-CoV-2 and the recCA of SARS-CoV-1 sharing 98.6% genetic identity with SARS-CoV-1.

### Phylogeography of bat sarbecoviruses related to SARS-CoV-1 and SARS-CoV-2

The phylogenies of the sarbecoviruses have an appreciable degree of geographic structuring linked to their hosts’ ranges (Fig. 3)^6^. To infer the spatiotemporal spread of the SARS-CoV-1-like and SARS-CoV-2-like viruses, we simultaneously fitted a spatially-explicit phylogeographic model to each NRR. As the sampling locations of SARS-CoV-1, SARS-CoV-2, and the pangolin sarbecoviruses likely do not represent where their direct bat virus ancestors circulated^16–19^, we excluded their locations from the primary phylogeographic analyses to avoid the impact of dispersal in non-bat hosts (Extended Data Fig. 2).

**Figure 3.**
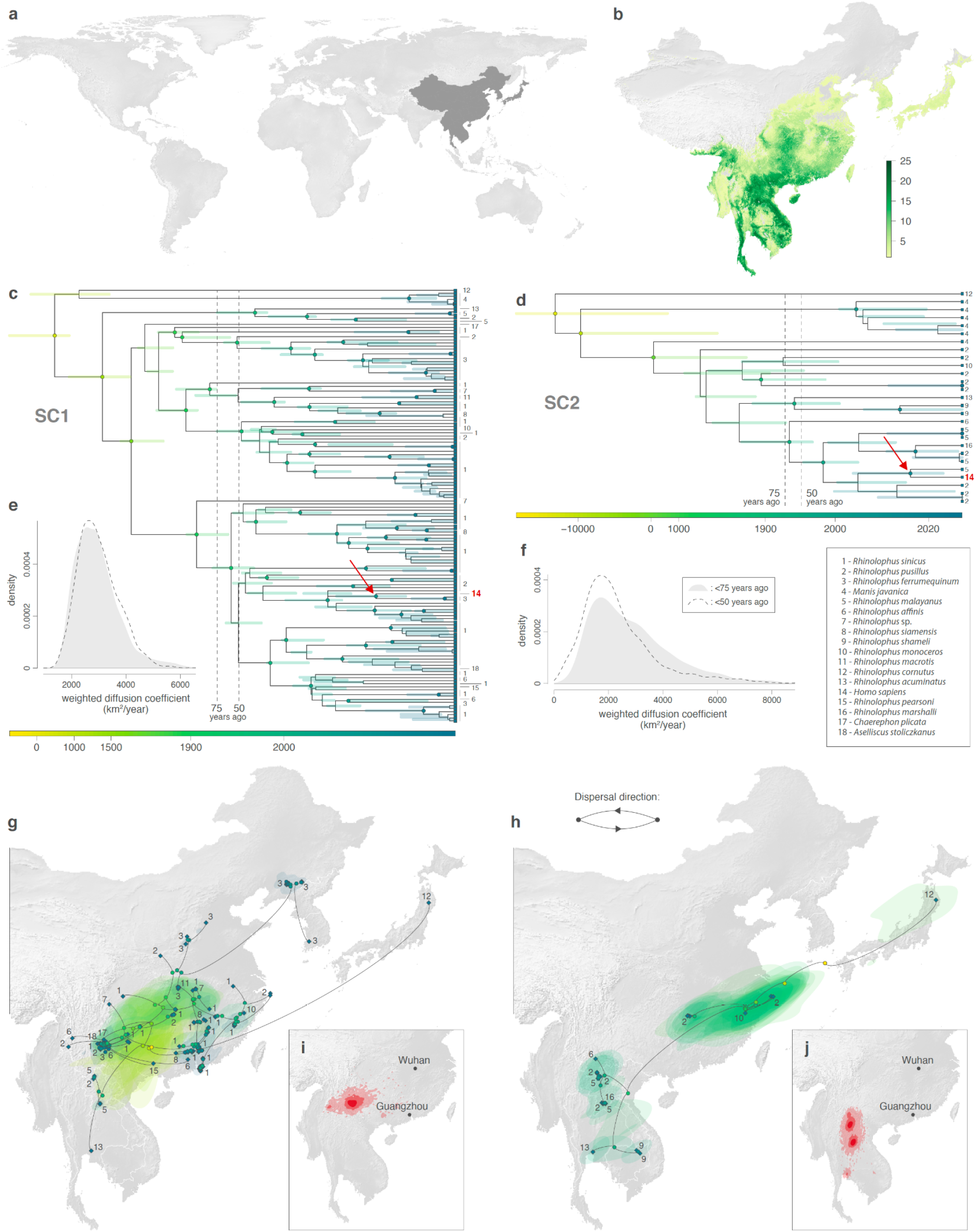
Estimated evolutionary history of SARS-CoV-1-like (SC1, on the left) and SARS-CoV-2-like (SC2, on the right) viruses. **a,** Global map with the study area highlighted in dark gray. **b,** Rhinolophid (horseshoe bat) species richness with the scale bar showing the number of co-occurring Rhinolophid species per km^2^. **c-d,** time-scaled maximum clade credibility (MCC) phylogenies obtained after correction for substitution saturation, shown on a log-transformed time scale. Horizontal segments displayed at each internal node indicate the 95% HPD interval of the estimated node age. Closest-inferred ancestor indicated by a red arrow. **e-f,** weighted diffusion coefficient estimates based on phylogenetic branches occurring less than 75 (plain light gray curves) and 50 (dashed curves) years ago. **g-h,** reconstruction of the dispersal history of viral lineages for two NRRs (NRR14 for SARS-CoV-1-like viruses, and NRR3 for SARS-CoV-2-like viruses) obtained through continuous phylogeographic inference (see Extended Data Fig. 9-12 for all other NRRs). Nodes are coloured from yellow (oldest most recent common ancestor) to blue (most recent collection date) and are superimposed on 80% HPD polygons coloured according to the same log-transformed time scale, reflecting the uncertainty of the Bayesian phylogeographic inference. 80% HPD polygons were only computed and reported for the last 1,000 years. Dispersal direction (anti-clockwise) of viral lineages is indicated by the edge curvature. In panels c-d-g-h, tip nodes are displayed as squares (as opposed to dots for internal ones) and are associated with a number indicating the host species from which the virus was sampled. The host species are numbered by decreasing order of sampling frequency. **i-j,** estimated position of the closest-inferred bat virus ancestors of SARS-CoV-1 (**i**) and SARS-CoV-2 (**j**) based on the continuous phylogeographic reconstruction of all NRRs. We display the 95%, 75% and 50% HPD regions with an increasing red color darkness.

Due to substitution saturation, the deeper evolutionary history of rapidly evolving viruses, including sarbecoviruses, are underestimated using standard molecular clock approaches^8,11,20,21^. To account for the effects of substitution saturation on the SARS-CoV-1-like and SARS-CoV-2-like phylogenies when trying to understand the long-term dispersal patterns of sarbecoviruses, we applied the Prisoner of War (PoW) model^8^—a mechanistic molecular clock model that estimates divergence times while accounting for time-dependent evolutionary rates—to the posterior phylogenies with phylogeographic estimates scaled in units of genetic distance (Extended Data Fig. 1; see Methods). We find that substitution saturation primarily affected the inferred time of divergence events at least 50 years before the most recently sampled virus in our dataset, with younger internal nodes having relatively unchanged ages and indicating that our inferences of the time of the closest-inferred ancestors are unbiased (Extended Data Fig. 7). The PoW-transformed phylogenies had tMRCAs that were at least several hundred to tens of thousands of years older (Extended Data Fig. 8).

To investigate the dispersal dynamics and history of the two sets of viruses, we computed their associated lineage diffusion coefficients^22^ (Fig. 3e,f and Extended Data Fig. 2) and separately represented the PoW-transformed phylogeny of each NRR across Asia (Fig. 3g,h, Extended Data Fig. 9-12). Both sets of viruses appear to have a few long lineage dispersal events, but the number of these dispersals is likely an underestimate due to undersampling of sarbecoviruses from various regions in Asia. The inferred dispersal history of viral lineages looks consistent among NRRs (Fig. 3g,h, Extended Data Fig. 9-12): we observe a similar spatial distribution of the sets of viruses and their connections on the different NRR maps. Moreover, the phylogeographic patterns appear to be spatially structured, with phylogenetically similar viruses being generally sampled in geographically similar regions and almost no back and forth travel of viral lineages across relatively large regions of the study area. Our analyses reveal a moderate to strong correlation between the patristic distance—the sum of the branch lengths connecting two nodes on a tree—and great-circle geographic distance computed for each pair of bat virus samples: a Spearman correlation coefficient of 0.45 for SARS-CoV-1-like viruses (95% HPD: 0.36-0.59) and of 0.66 for SARS-CoV-2-like viruses (95% HPD: 0.12-0.83) (see Supplementary Table 5 for a comparison with other viruses).

Either the root (Fig. 3d) or a deep ancestral node (see Data and Code Availability) of the SARS-CoV-2-like phylogenies consistently corresponds to the split between Rc-o319, sampled in Japan in 2013, and virus relatives from China and Southeast Asia for most NRRs. Our PoW-calibrated analysis gives a mean date of 8259 BCE across all NRR tMRCA median heights for this node (range: 23,777 BCE–1381 CE), providing further support for the ancient origins of the SARS-CoV-2-like viruses and therefore sarbecoviruses as a whole^7,8^. Conversely, although most NRRs of each set of viruses had a tMRCA of at least several thousands of years old, NRRs 20 (Orf1b) and 31 (N, Orf10) of the SARS-CoV-1-like viruses had tMRCAs of 1429 CE (95% HPD: 970 CE–1642 CE) and 1730 CE (95% HPD: 1625 CE–1828 CE), respectively, suggesting that these NRRs experienced selective sweeps, reducing genomic diversity in these genomic regions in the relatively recent past.

### Location of the closest-inferred ancestors of SARS-CoV-1 and SARS-CoV-2

The PoW transformation shows that the timing of divergence events in the last 50 years inferred from standard molecular clock models are unbiased (Extended Data Fig. 7). However, as the PoW model enforces contemporaneous tips, it can produce unreliable results for the most recent inferred ages. Therefore, we performed a separate tip-dated phylogeographic analysis to infer where the closest-inferred ancestors of each SARS-CoV circulated in the past 50 years (Extended Data Fig. 2). We find that the closest-inferred ancestors of SARS-CoV-1 likely circulated in western China (i.e., Yunnan, Sichuan, or Guizhou). The closest-inferred ancestors of SARS-CoV-2 likely circulated in Yunnan, China or northern Laos, overlapping with a set of contiguous cave structures extending through these regions^23,24^. Our results indicate that both SARS-CoV-1 and SARS-CoV-2 emerged in humans over a thousand kilometers from where their closest-inferred bat virus ancestors likely circulated. The inferred geographic location of the closest-inferred ancestors of each SARS-CoV additionally overlapped with regions with moderate Rhinolophid (horseshoe bat) species richness (Fig. 3b and Extended Data Fig. 13–15), including in regions known to harbor *R. affinis* and *R. ferrumequinum* (Extended Data Fig. 13), and the closest-inferred ancestor of SARS-CoV-2 additionally overlapped with known locations of *R. malayanus* circulation (Extended Data Fig. 14). Including the location of the human and pangolin SARS-CoVs as a robustness analysis resulted in the inferred regions where the closest-inferred ancestors of both SARS-CoV-1 and SARS-CoV-2 circulated to include where these viruses also emerged (Extended Data Fig. 16, 17); however, these inferences are likely biased by the non-bat travel (*e.g.* intermediate hosts associated with wild and farmed animal supply chains) of the ancestors of SARS-CoV-1 and SARS-CoV-2 and trafficking^18^ of the pangolin hosts.

### Sarbecoviruses spread at rates similar to their bat hosts

To investigate how fast the SARS-CoV-1-like and SARS-CoV-2-like viruses recently spread across Asia, we inferred the weighted diffusion coefficient, a measure of the diffusivity of viral transmission, for the 50 years before the most recently sampled genome in each dataset using our tip-dated phylogenies (Extended Data Fig. 2). We found that the SARS-CoV-1-like viruses spread with a weighted diffusion coefficient of 2884 km^2^/yr (95% HPD: 1654–4358; Fig. 3e), and the SARS-CoV-2-like viruses circulated with a weighted diffusion coefficient of 2050 km^2^/yr (95% HPD: 233–5470; Fig. 3f). The SARS-CoV-1-like viruses were likely inferred to spread, on average, slightly more quickly than the SARS-CoV-2-like viruses because of the recent dispersal events to South Korea. These results were robust to the inclusion of the locations of the human and pangolin SARS-CoVs and were similar to the diffusion coefficients calculated for the 75 years preceding the most recently sampled genomes (Supplementary Table 5). Horseshoe bats, the primary hosts of sarbecoviruses, tend to forage within around 2-3 km of their roost^25–28^ and have a reported diffusion coefficient of 1999 km^2^/yr^29^. Therefore, both the SARS-CoV-1-like and SARS-CoV-2-like viruses (in the context of host infections) traveled at a similar rate as their hosts, suggesting that these viruses are able to efficiently explore new niches, likely due to the density of Rhinolophid roosts.

### Direct ancestors of the SARS-CoVs likely could not have reached sites of emergence via the bat reservoir alone

To understand whether the bat virus ancestors of each SARS-CoV could have spread to where the SARS-CoVs emerged in humans via dispersal of SARS-related coronaviruses across bat populations, we determined the minimum distance from the location of the closest-inferred bat virus ancestor to the province of emergence (Fig. 3i,j), and given the time between emergence and this ancestor (Fig. 1b,c), we calculated the dispersal velocity this implies. We compare this rate to the dispersal velocity of bat virus lineages to obtain a posterior rank distribution (Extended Data Fig. 18). For the closest-inferred bat virus ancestor of SARS-CoV-1 to have reached Guangdong Province, it would have needed a dispersal velocity greater than 95% of the dispersal velocities of bat virus lineages across 8 NRRs, including NRR 14 (Extended Data Fig. 18a). Similarly, for the direct bat virus ancestor of SARS-CoV-2 to have reached Hubei Province, it would have needed a dispersal velocity greater than 95% of the dispersal velocities of bat virus lineages across 7 NRRs, including NRRs 3 and 6 (Extended Data Fig. 18b) and NRRs with a time of the closest-inferred ancestor as early as 2009 (Fig. 1c). Considering the large distance separating the closest-inferred bat virus ancestors from where SARS-CoV-1 and SARS-CoV-2 each emerged (Fig. 3i,j) and the high dispersal velocities necessary to traverse the aforementioned distance (Extended Data Fig. 18), it is very unlikely that the lineages descending from the closest-inferred bat virus ancestors of SARS-CoV-1 and SARS-CoV-2 reached, respectively, Guangdong Province and Hubei Province solely via dispersal of these viruses through their bat reservoirs.

## Discussion

Our study reveals viral genomic sequence fragments closely related to the very recent ancestors of SARS-CoV-1 and SARS-CoV-2 are present in published genomes sampled from Rhinolophid (horseshoe) bats. By reconstructing the evolutionary history of the non-recombinant fragments of sarbecoviruses, we show that the ancestors of SARS-CoV-1 and SARS-CoV-2 circulated in bat populations hundreds to thousands of kilometers away from the sites of, and as recently as 1–3 years prior to, the emergence of these viruses in humans.

Sarbecoviruses undergo extensive recombination and tend to share breakpoints at specific genome regions, so-called recombination hotspots^4,30^. Although whole-genome comparisons previously suggested there may have been decades of separation between the closest-inferred bat virus ancestor and each of the SARS-CoVs^31^, the recency of the closest-inferred ancestor in multiple subgenomic regions indicate that whole-genome comparisons are an unreliable measure of evolutionary divergence. It is, therefore, the non-recombinant fragments that must be analyzed to properly understand the emergence of the human SARS-CoVs^5,6^. We additionally show that adding new genomes to the recombination-aware ancestor reconstruction brought the sequence identity of the recombinant common ancestor of each SARS-CoV to its respective human virus, closer to 100%. This was true even though the new genomes did not always share a higher whole-genome identity to the SARS-CoVs than previously published sarbecovirus genomes.

The phylogeographic approach implemented here provides detailed insights into the sarbecoviruses’ extensive historical movement across China and Southeast Asia. We find that the viral lineages diffuse at a similar rate to their primary hosts: small horseshoe bats. These bats typically have small home ranges (2-3km^2^ per night), are rarely migratory^27,32^, and have a limited dispersal capacity with scant evidence of longer migrations (and limited to species in temperate rather than tropical regions)^33,34^. After accounting for recombination, the closest-inferred bat virus ancestors of SARS-CoV-1 and SARS-CoV-2 are inferred to have circulated in regions spanning from Northern Laos to Western China.

Considering that the closest-inferred bat virus ancestors circulated in these regions almost immediately preceding the emergence of SARS-CoV-1 and SARS-CoV-2, our results suggest that there would not have been enough time for the direct bat virus ancestor to reach the location of emergence of the human SARS-CoVs via normal dispersal through bat populations alone. SARS-CoV-1 is thought to have been moved from bats in Yunnan province to Guangdong province via intermediate mammalian hosts^16^. The movement of palm civets and raccoon dogs as part of the wild and farmed animal trade has been implicated in the emergence of SARS-CoV-1^35^, suggesting that human-mediated transport of these animals can allow for viral lineages to move faster than their bat hosts (Extended Data Fig. 18). Given (i) the presumed emergence of SARS-CoV-1 through the animal trade, (ii) the recency and location of the closest-inferred ancestor relative to SARS-CoV-2, and (iii) the clear evidence that the epicenter of the SARS-CoV-2 pandemic at one of only four markets in Wuhan that sold live wildlife from plausible intermediate host mammal species^36,37^, the direct ancestor of SARS-CoV-2 likely moved from an area in or around Yunnan province, to Hubei province, via the wild and farmed animal trade.

The similar rates of diffusion among viral lineages and horseshoe bats suggest these viruses are able to rapidly and efficiently explore their environmental niches. The sarbecoviruses are therefore being transmitted among dense bat populations of the same and different species, as is also evidenced by the high levels of recombination in bat sarbecoviruses. For example, viral movement is likely to take place through horseshoe bats regularly moving among roosts, especially as roost switching tends to happen during breeding season or if bats travel or migrate to hibernacula^28,38^. Furthermore, in hibernacula horseshoe bats will roost in very dense aggregations with different species^39^, providing the potential for spillover between bat species whilst bodily functions are downregulated and body temperatures can reach very low levels^39,40^. Heterothermy during hibernation (and in some species even sleep) and variable temperatures among species may also impact arousal patterns and vulnerability to pathogens^41,42^.

Despite limited evidence for long-distance travel of horseshoe bats, there are several relatively long-distance viral lineage dispersals between nodes in the phylogenies. These geographically distant nodes therefore likely represent a lack of sarbecovirus sampling from areas between the distant locations rather than long-range bat movements, though it could also indicate dispersal from more migratory conspecific bat species, such as *Chaerephon* which can share roosts. For example, internal branches of the SARS-CoV-2-like tree represent movement from the east of China (Zhejiang) to the west (Yunnan), to Thailand and Cambodia (Fig. 3h). There are likely many more, still-undiscovered SARS-CoV-2-like virus populations that would bridge the geographic gap between these regions (e.g., in northern Vietnam, Guizhou, and Guangxi) as diversity of bats remains high across these regions (Figure 3i).

By accommodating time-dependent rate variation in our phylogeography analysis, we are able to infer the deeper evolutionary history of sarbecoviruses more accurately than previous efforts. For example, one ancestral node connecting distant regions is that between the Chinese and Southeast Asian SARS-CoV-2-like viruses and the Japanese sarbecoviruses (Rc-o319 represented in our dataset, but more viruses similar to SARS-CoV-2 have been recently sampled in Japan^43^), and we infer an average estimate of 8,259 BCE for this node (the oldest median estimate—NRR 21—being earlier than 20,000 years BCE). Rhinolophid bats will not fly over the open sea deliberately^44–47^, but can be blown over narrow ocean channels or travel on ships^48^. However, during the Last Glacial Maximum (LGM), until approximately 8,000 BCE, Japan was likely connected to the mainland through land bridges both in the north (connecting Hokkaido to Siberia) and south (connecting Kyushu to China)^49^. This most recent time when Japan was connected to the mainland aligns well with our dating estimates of when the Japanese bat sarbecovirus clade separated from the mainland clade. Additionally, there is evidence of mammals moving through the land bridges during the LGM^50^. The bat species endemic to Japan known to harbour sarbecoviruses, *R. cornutus*, and its phylogenetically closest mainland species infected by SARS-CoV-2-like viruses, *R. pusillus*, are estimated to have shared a common ancestor 13.3 mya^51^. This much more ancient separation between the host species indicates that the viruses did not co-evolve with the bats. Instead, mainland bats likely transmitted the ancestral SARS-CoV-2-like viruses to the Japanese bat populations through movement via land bridges during the LGM.

Whilst horseshoe bat ranges extend across most of China, diversity decreases at increasing latitudes, and virus sampling in bats has largely been limited to higher diversity areas, including Yunnan province in China and Laos. Additionally, other large areas with high diversity, such as Northern Vietnam, remain unsampled and host a similar community of bat species. In the absence of bat viruses sampled closer to where SARS-CoV-1 and SARS-CoV-2 emerged, the inference of the location of the closest-inferred bat virus ancestors is biased toward where published, closely related viruses were sampled. The SARS-CoV-1-like bat viruses are better represented (n=134) and therefore have more phylogeographic resolution than the SARS-CoV-2-like bat viruses (n=20). Although there are viruses closely related to the SARS-CoVs across most of the NRRs, the viruses are relatively more divergent across the NRRs that overlap with the Spike gene (Fig. 1, Extended Data Fig. 6). This relatively greater divergence suggests that Spike genes that can bind non-bat entry receptors circulate in lower frequencies in wild horseshoe bat populations and, therefore, are less likely to be sampled. Further sampling of closely related sarbecoviruses, methodological improvements in disentangling complex recombination patterns, and improved molecular clock calibration can allow for the inference of more detailed spatiotemporal trends and recombination patterns, as well as more complete evolutionary histories of each NRR.

Since the emergence of SARS-CoV-1 and SARS-CoV-2 in humans, the descendants of the bat viruses that gave rise to them have experienced years of evolution and almost certainly recombined with yet-unsampled lineages of the sarbecovirus tree. Hence, we should not expect to find the direct ancestor of SARS-CoV-1 or SARS-CoV-2 circulating in wild bats in future sampling. The search for, and analyses of, sarbecoviruses related to SARS-CoV-1 and SARS-CoV-2 have often focused on partial genomes, typically the RdRp protein^52–57^, but future sequencing efforts should aim for whole genome sequences to detect and characterize all genomic fragments that descend from closely related ancestors. The non-recombinant segments of these mosaic genomes are pieces of a complex puzzle that hold the key to understanding the intricate mechanisms behind the emergence of past and future SARS-CoVs into the human population.

## Supporting information

Sarbecovirus genomes metadata.

Genome accessions.

## Materials and Methods

### Dataset compilation

For the animal sarbecovirus dataset we compiled a set of 167 whole-genome sequences of the subgenus *Sarbecovirus* (available as of March 2022) sampled primarily in horseshoe bat hosts, including reference genomes of the human viruses SARS-CoV-1 and SARS-CoV-2 and six pangolin-infecting viruses. All sequences were publicly available as of October 2022 (Supplementary Data File 1). Collection date, sampling location and host metadata associated with the sequences were retrieved from the corresponding sequence databases and manually checked with the published papers describing the sequences. For sequences where exact city/cave sampling location information was provided a single longitude and latitude data point corresponding to that location was recorded. For cases where only province-specific sampling information were available a six point geographic location range encompassing each province was recorded. The metadata for these 167 genomes are available in Supplementary Data File 1.

For the SARS-CoV-2 dataset, we used the 787 genomes and sampled 1,000 trees from the posterior tree distribution from Pekar et al.^15^ For the SARS-CoV-1 dataset, we downloaded the 98 publicly available whole genomes of SARS-CoV-1 from humans and civets from Genbank, with dates and locations matched by metadata or literature review: https://github.com/andersen-lab/SARS-CoV-1_Evolution/blob/main/metadata_extended.csv. A list of all accessions used in this study are available in Supplementary Data File 2. All GISAID accessions are also available through GISAID when using the identifier *EPI_SET_230410yb*.

### Recombination analysis

The 167 whole-genome sequences were aligned using MAFFT (v7.453, localpair option) ^58^ and manually curated using BioEdit^59^. To explore the recombination patterns of SARS-CoV-1-like and SARS-CoV-2-like sarbecoviruses separately we selected a number of representatives from each set of genomes on which to perform recombination analysis using the Genetic Algorithm for Recombination Detection (GARD)^13^. The genome selection was made by first performing a preliminary phylogenetic analysis of the relatedness between these genomes using the recombination breakpoints presented in Lytras et al. (2022)^4^. Based on the preliminary analysis, clusters of genomes grouping with SARS-CoV-1 and SARS-CoV-2 for at least one of their genomic segments and showing the same overall recombination pattern were manually identified and one representative genome was selected from each cluster for the corresponding downstream GARD analyses. For the SARS-CoV-2-like set, 19 genomes including the SARS-CoV-2 reference Wuhan-Hu-1 were used in the GARD analysis. For the SARS-CoV-1-like set, 38 genomes including the SARS-CoV-1 HSZ-Cc genome were used in the analysis (Supplementary Data File 1). The representative sequences for each set were retrieved from the full alignment and the extended 3’ and 5’ ends were trimmed from each resulting subset alignment to reduce sequence noise in the recombination analysis. GARD was performed for each subset alignment using the HyPhy software suite v.2.5.29^60^ under a general time reversible (GTR) substitution model and a 3-class general discrete distribution (GDD) to account for site-to-site rate variation. The SARS-CoV-2-like GARD analysis finished with a total of 72 consecutive breakpoint models, while the SARS-CoV-1-like analysis finished with 56. Based on these models, we accepted positions as confident recombination breakpoints if they were present in at least 1/3 of the consecutive models and breakpoints closer than 100 alignment positions to one another were merged. The resulting breakpoint positions were mapped back to the full whole-genome alignment which was subsequently split into the corresponding non-recombinant regions (NRR) for each set of breakpoints. To avoid uninformative alignment regions biasing phylogenetic inferences, all NRR alignments were processed using HMMCleaner^61^ before performing any downstream analyses. The GARD breakpoint results for the SARS-CoV-2-like set have previously been used in Martin *et al.* (2022)^62^.

To separate all genomes into SARS-CoV-1-like and SARS-CoV-2-like viruses we first inferred maximum likelihood phylogenies for every NRR alignment using IQ-TREE (v.1.4.4)^63^ under a GTR+I+F+G4 substitution model, assessing node support by performing 10,000 replicates of the ultrafast bootstrap approximation for each inference^64^. All trees were rooted by the outgroup clade of sarbecovirus genomes sampled in Europe and Africa (accessions: NC_014470, MZ190137, KY352407, MT726043, MT726044, MT726045, MZ190138, MW719567) and nodes with ultrafast bootstrap support below 70 were collapsed using the ‘ete3’ python package^65^. Based on the resulting topologies, the viruses clustering monophyletically with SARS-CoV-2 but not with SARS-CoV-1 for at least one of the SARS-CoV-2-like GARD NRR trees were placed in the SARS-CoV-2-like viruses and vice versa for the SARS-CoV-1-like virus set. This led to two non-independent sets of 27 SARS-CoV-2-like and 140 SARS-CoV-1-like genomes (each including SARS-CoV-2 and SARS-CoV-1 respectively) used in the downstream dating and phylogeographic analyses presented in the paper.

### Comparison of breakpoint positions

Based on the GARD recombination analysis, 30 and 26 breakpoint positions were detected for the two virus subgroups. Only one genome was used in both GARD sets (Supplementary Data File 1), so overall breakpoint inferences are expected to represent independent recombination patterns of known SARS-CoV-1-like and SARS-CoV-2-like viruses respectively. We calculated the distances between the closest position pairs between the two breakpoint sets, resulting in 26 pair distances (each SARS-CoV-2-like position matched with its closest SARS-CoV-1-like position). To compare whether the pairs were more or less closely located than one would expect by chance, we performed 100 simulations of position sets randomly sampled across the same alignment length without replacement (one set with 26 positions and one with 30 for each simulation to match the empirical data on the number of breakpoints). We then quantified the distances between closest pairs in the simulated sets as done with the observed breakpoints. Both empirical and simulated distances formed distributions with log-normal shapes, hence 90% confidence intervals and means were estimated using Cox’s method for inferring log-normal confidence intervals^66^. Similarly, the significantly closest observed breakpoint pairs were defined as those with distances below the one-tailed lower 90% confidence interval of the observed log-normal distance distribution.

### Molecular clock calibration

There is insufficient data to inform a molecular clock when using tip dating with sarbecoviruses sampled from bats and pangolins^6^. To identify a suitable rate prior for each NRR identified from the recombination analysis, we conducted a phylogenetic analysis of SARS-CoVs sampled from humans.

Molecular clock calibration across the 27 inferred NRRs of the SARS-CoV-2-like viruses was conducted using a Bayesian approach with BEAST v1.10.5^67^. We inferred the substitution rate of SARS-CoV-2 across each of these NRRs by using an empirical tree distribution (n=1,000) from our previously published Bayesian phylogenetic analysis of 787 early pandemic genomes (Supplementary Data File 2), sharing the topologies and substitution models across all the NRRs, but assuming an independent and estimable strict clock rate under an uninformative continuous-time Markov chain (CTMC) rate prior^68^ for each NRR. We ran one Markov chain Monte Carlo (MCMC) chain of one million generations, subsampling every one thousand iterations to a continuous parameter log file. The first 15% of the chains were discarded as burn-in. Convergence and mixing was assessed in Tracer v1.7.1^69^ and all relevant effective sample size (ESS) values were >200.

To calibrate the molecular clock for the SARS-CoV-1-like viruses analysis, we first performed Bayesian phylogenetic inference on 98 human and civet SARS-CoV-1 genomes (Supplementary Data File 2), using an HKY85 substitution model with a gamma site heterogeneity model and 4 gamma categories (G4), a strict molecular clock, and an exponential growth coalescent prior. We ran a single chain of 100 million generations, subsampling every 10 thousand iterations to a continuous parameter log and tree files. We next inferred the substitution rate of SARS-CoV-1 across the 31 inferred NRRs of the SARS-CoV-1-like viruses by using an empirical tree distribution (n=1,000) from these results, sharing the topologies and substitution models across all the NRRs, with independent and estimable clock rates under CTMC rate priors. We ran one chain of one million generations, subsampling every one hundred iterations to a continuous parameter log file. For each step of the inference, the first 15% of the chains were discarded as burn-in. Convergence and mixing was assessed in Tracer and all relevant effective sample size (ESS) values were >200.

### Bayesian divergence time estimation

We used BEAST to perform phylogenetic inference and ancestral state reconstruction across the 27 NRRs of the SARS-CoV-2-like viruses. For the primary analysis (Extended Data Fig. 2), we used all of the 27 SARS-CoV-2-like genomes (including SARS-CoV-2), an uncorrelated relaxed clock using calibrated region-specific normally-distributed substitution rate priors for each NRR based on SARS-CoV-2 sampled from humans, a GTR+G4 substitution model, a non-parametric skygrid prior^70^ with 49 grid points and a cutoff of 1,000 (which translates to 1022 CE), and Hamiltonian Monte Carlo gradient-based sampling for node ages and branch rates^71,72^. Specifically, we set the mean and standard deviation of the rate prior for each NNR to equal the posterior mean and standard deviation estimated during molecular clock calibration with SARS-CoV-2 genomes. We tracked the stem age of SARS-CoV-2 (*i.e.*, where SARS-CoV-2 connects to the rest of the phylogeny, also referred to as the time of the closest-inferred ancestor, with the parent node of SARS-CoV-2 referred to as the closest-inferred ancestor). For each NRR, we ran one chain of 10 million generations, subsampling every one thousand iterations to continuous parameter log and tree files. The first 15% of the chain was discarded as burn-in for most of the analyses; for several analyses we had to discard a greater percentage of the chain to account for poor initial mixing. Convergence and mixing was assessed in Tracer, and all relevant ESS values were >200.

We performed the same analysis for the SARS-CoV-1-like viruses for its 31 NRRs, using all of the 140 SARS-CoV-1-like genomes (including SARS-CoV-1). The BEAST parameterization was identical to the analysis for the SARS-CoV-2-like viruses, except we (1) tracked the stem age of SARS-CoV-1 (*i.e.*, the time of the closest-inferred ancestor of SARS-CoV-1), (2) used the relevant calibrated molecular clock priors for the 31 NRRs of the SARS-CoV-1-like viruses, and (3) ran one chain of 100 million generations, sub-sampling every 10 thousand iterations to continuous parameter log and tree files.

### Bayesian divergence time estimation sensitivity analyses

We performed a series of sensitivity analyses for the Bayesian divergence time estimation: (i) dividing the standard deviation of the molecular clock prior by 5, (ii) dividing the standard deviation of the molecular clock prior by 10, and (iii) including an additional human SARS-CoV-1 genome and a civet SARS-CoV-1 genome in the SARS-CoV-1-like viruses analyses. For (iii), SZ3 (AY304486) was selected as representative of the earliest sampled detection in the intermediate animal reservoir (market-based civets). HZS2-C (AY394992) was selected to represent the early stages of the human epidemic from late January 2003 onwards.

### Constructing the recombinant common ancestor

For each NRR of the SARS-CoV-2-like viruses, we reconstructed the most likely nucleotide sequence belonging to the parent node of SARS-CoV-2 based on the tree distribution from the primary Bayesian divergence time estimation (Extended Data Fig. 2). The most likely nucleotide sequences for the 27 NRRs are then concatenated to form the recombinant common ancestor (recCA) of SARS-CoV-2.

To analyze how the recCA changed as more genomes were released, we constructed the recCA of SARS-CoV-2 as the genomes of closely related sarbecoviruses were chronologically published. We performed ancestral state reconstruction using the posterior trees with all 27 taxa. We then identified the inferred sequence of the most recent common ancestor of SARS-CoV-2 and its most closely related taxa based on the published genomes as new genomes were sequenced. We then examined the genetic similarity of the recCA, as well as the most closely related published genome, to SARS-CoV-2 as a function of the publication date. The same analysis was performed for SARS-CoV-1 and the SARS-CoV-1-like viruses.

### Bayesian phylogeographic inference

We inferred the evolutionary and geographic history of the 27 NRRs of the SARS-CoV-2-like viruses with BEAST and using the BEAGLE library v3 to increase computational performance^73^. In order to optimally inform a flexible phylogeographic reconstruction approach, we jointly fit a bivariate continuous diffusion model using a shared estimable precision matrix across all independent random NRRs phylogenies, rescaling this matrix across branches in each tree to model a relaxed random walk (RRW)^74^. All branch-specific RRW rate scalars are drawn from the same log-normal distribution with a mean of 1 and an estimable standard deviation. In the continuous phylogeographic diffusion model, we exclude the sampling locations for the human and pangolin viruses and integrate over the uncertainty for locations that lack precision. As a tree prior, we specify a joint non-parametric skygrid prior with 49 grid points and a cutoff of 1,000 (which translates to 1022 CE) or a cutoff of 0.5 substitutions per site for trees scaled in units of genetic distance (see below). We further specify a joint HKY85 nucleotide substitution model across NRRs and independent strict molecular clock models; these relatively simple sequence evolutionary parameterizations were chosen to meet the requirements for the Prisoner of War (PoW) model^8^ to rescale the evolutionary histories (see below). We perform two versions of this analysis (Extended Data Fig. 2): one with the clock rate priors also used in the previously described divergence time estimation to estimate the trees in units of time and one with clock rates fixed to 1 and contemporaneous tips in order to estimate trees in substitution units for PoW transformation. We ran two chains of 500 million generations, subsampling every 10 thousand iterations to continuous parameter log and tree files; the first 10% of the chain was discarded as burn-in. Convergence and mixing was assessed in Tracer, and all relevant ESS values were >200.

We performed the same analysis for the SARS-CoV-1-like virus set for its 31 NRRs. The BEAST parameterization was identical to the analysis for the SARS-CoV-2-like viruses, but here we use the rate priors with standard deviations divided by 5. We ran 10 chains for > 200 million generations, subsampling every 500 thousand iterations to continuous parameter log and tree files; the first 10% of the chain was discarded as burn-in. Convergence and mixing was assessed in Tracer, and all relevant ESS values were >200.

To account for the time-dependent decline in the evolutionary rate estimates of sarbecoviruses, we applied the PoW model, a mechanistic evolutionary method for estimating virus substitution rates over long evolutionary timescales. For this, we used the post-burn-in substitution trees inferred from the previous step along with the posterior rate distributions for each NRR of the SARS-CoV-1-like viruses and the SARS-CoV-2-like viruses to rescale substitution trees into time trees using the PoW transformation (see ref Ghafari et al.^8^ for more details on the method).

We used TreeAnnotator 1.10^67^ to obtain and annotate a maximum clade credibility (MCC) tree for each continuous phylogeographic reconstruction. Finally, we used the R package ‘seraphim’^75^ to extract the spatiotemporal information embedded within 100 trees sampled from each post-burn-in posterior distribution and to visualize the different phylogeographic reconstructions. For each NRR, we estimated the weighted diffusion coefficients^22^ with the ‘spreadStatistics’ function of the R package ‘seraphim’^75^.

As a sensitivity analysis, we repeated these phylogeographic analyses using the sampled locations of the pangolin and human viruses.

### Ranking the dispersal velocity of the lineage descending from the closest-inferred ancestor

Using the posterior tree distribution from the tip-dated phylogeography that excluded the human and pangolin viruses, we calculated the dispersal velocity across each branch of each post-burn-in posterior tree in the 50 years preceding the most recent tip, excluding the branches leading to the human or pangolin viruses (or their clades, if the clade was entirely composed of pangolin and human viruses). These dispersal velocities were pooled for each NRR, resulting in a dispersal velocity distribution per NRR. We next computed the minimum distance between the closest-inferred ancestor and Guangdong province or Hubei province for SARS-CoV-1 or SARS-CoV-2, respectively. The necessary dispersal velocity for the closest-inferred ancestor to reach either Guangdong or Hubei province was calculated using this minimum distance and the branch leading to, respectively, SARS-CoV-1 or SARS-CoV-2 (*i.e.*, the time between closest-inferred ancestor and the SARS-CoV). We then computed the rank of this dispersal velocity by comparing it against the dispersal velocity distribution of the given NRR, resulting in a posterior distribution of dispersal velocity ranks for the lineages descending from the closest-inferred ancestors of SARS-CoV-1 and SARS-CoV-2 to reach their provinces of emergence.

### Modeling of bat distributions

Our bat distribution database comprised of the data used in recently published studies^76^, recent Global Diversity Information Facility (GBIF) data^77,78^, and the Darkcide database^79^. The Darkcide database included recent data collected by bioacoustic surveys in Chiang Mai and Kanchanaburi in Thailand by A.C. Hughes using an echometer Pro, and multiple published datasets^76–78,80–88^. The data was integrated as we attempt to keep the database complete, even if not all data is used within our particular study. All Rhinolophid species, as well as *Chaerephon plicatus* and *Aselliscus stoliczkanus* were extracted from the dataset, and then the dataset was clipped to Southeast Asia (India to the West, Mongolia to the North, Japan to the East and Malaysia to the South) to buffer the region of interest with distribution data. Once all duplicate records had been removed, the clipped data for Rhinolophids included 10,254 distribution points for all species across the region, in addition to 235 records for *A. stoliczkanus* and 108 for *C. plicatus*.

For environmental data, we based variable selection on those used in Zhou et al. (2021)^76^ and known from our previous work to be important to bats in the region^89,90^. All analysis was conducted in ArcMap 10.8 (ESRI) at a resolution of 0.008°, which is approximately 1km^2^, and all layers were clipped and subsampled to the same resolution using a mask. Climate variables were downloaded from Chelsa^91^ (available at https://chelsa-climate.org/downloads/) based on a selection from our previous analyses (the bioclimatic variables: Annual Mean Temperature, Mean Diurnal temperature Range, Isothermality, Temperature Seasonality, Maximum and Minimum Temperature, Annual Precipitation, Precipitation seasonality, Maximum and Minimum precipitation)^89,90^. From Envirem (https://envirem.github.io/) we used the Thornthwaite Aridity index, climate moisture index, continentality, embergers pluviothermic quotient, growing degree days above 0, and both annual mean potential evapotranspiration and seasonality of potential evapotranspiration. In addition we used actual evapotranspiration^92^ (Trabusco & Zomer 2019), a vegetation canopy height layer^93^, forest canopy height^94^, tree density^95^, distance to bedrock^96^ (the bdticm and bdricm aggregated grids available at https://github.com/ISRICWorldSoil/SoilGrids250m/), and bulk carbon stock ^96^. These were all trimmed to the same region that bat data was clipped to and then species distributions were modelled using Maxent^97^ using the GUI following the same approach as Hughes et al. (2012)^89^. For each species, three replicates were run and the average was used. These probabilistic distribution maps were then reclassified using the 10th percentile cumulative logistic threshold to produce a binary presence-absence map with ArcMap 10.8, and all models had an AUC exceeding 0.85. The models were then clipped to the study region, and any Rhinolophids with no modelled suitable habitat in the clipped study area were removed from analysis. As the projections show suitable habitat but do not account for species biogeography we then cross-checked the 40 remaining Rhinolophid maps against the IUCN database (https://www.iucnredlist.org/), and species not listed were checked against relevant publications on the species. Based on the IUCN mapped ranges species endemic to Japan, or species not found in Japan or at similar latitudes (areas of the South Korean Peninsula are climatically suitable for a number of species, but lack continuous habitat to reach these regions if largely distributed in Southeast Asia) were then clipped to the biogeographically suitable range. The binary maps were then stacked using the Mosaic to New Raster function in ArcMap 10.8 to find the maximum number of species that may be co-distributed.

## Data and Code Availability

All genome accessions are available in Supplementary Data File 2. The XML files, code, and newick trees are available at https://github.com/phylogeography/sarbecovirus_phylogeography.

## Acknowledgements

S.D. acknowledges support from the *Fonds National de la Recherche Scientifique* (F.R.S.-FNRS, Belgium; grant n°F.4515.22) and from the Research Foundation - Flanders (‘Fonds voor Wetenschappelijk Onderzoek - Vlaanderen’, FWO, Belgium; grant n°G098321N). S.D. and P.L. acknowledge support from the European Union Horizon 2020 project MOOD (grant agreement n°874850). P.L., A.M. and M.A.S. acknowledge support from the European Union’s Horizon 2020 research and innovation programme (grant agreement no. 725422-ReservoirDOCS), from the Wellcome Trust through project 206298/Z/17/Z and from the National Institutes of Health grants R01 AI153044, R01 AI162611 and U19 AI135995. P.L. also acknowledges support from the Research Foundation - Flanders (‘Fonds voor Wetenschappelijk Onderzoek - Vlaanderen’, G0D5117N and G051322N). T.I.V. acknowledges support from the Branco Weiss Fellowship. D.L.R. and J.H. acknowledge funding from the UK Medical Research Council (MRC, MC_UU_12014/12, MC_UU_00034/5 and MR/V01157X/1). S.L. is funded by an MRC studentship. M.G. is supported by the Biotechnology and Biological Science Research Council (BBSRC) grant number BB/M011224/1. J.E.P. acknowledges support from NIH (T15LM011271) and the UC San Diego Merkin Fellowship. J.O.W. acknowledges support from NIH (R01AI135992). We gratefully acknowledge the authors from the originating laboratories and the submitting laboratories, who generated and shared through GISAID the viral genomic sequences and metadata on which this research is based (Supplementary Data File 2). We thank Kristian Andersen, Alexander Crits-Christoph, Fabian Leendertz, Niema Moshiri, Richard Neher, and Andrew Rambaut for fruitful conversations and insight. We gratefully acknowledge support from Advanced Micro Devices, Inc. with the donation of parallel computing resources used for this research.

## Author contributions

J.E.P., S.L., D.L.R., M.W., J.O.W., and P.L. conceived and planned the research. J.E.P., S.L., S.D., M.G., A.M., E.P., and P.L. analyzed and processed the data. A.M., T.I.V., M.A.S., J.H., S.D., J.O.W., and P.L. advised on statistical methodology. J.E.P., S.L., S.D., M.G., and P.L. designed and implemented genomic data processing pipelines. A.C.H modelled bat distributions. J.E.P., S.L., S.D., and P.L. wrote the first draft of the manuscript. All authors contributed to editing and interpreting the results. D.L.R., S.D., M.W., J.O.W, and P.L. jointly supervised the work.

## Corresponding and requests for materials

should be addressed to J.E.P., S.L., S.D]., M.W., J.O.W., or P.L.

## Extended Data Figures

**Extended Data Fig. 1.**
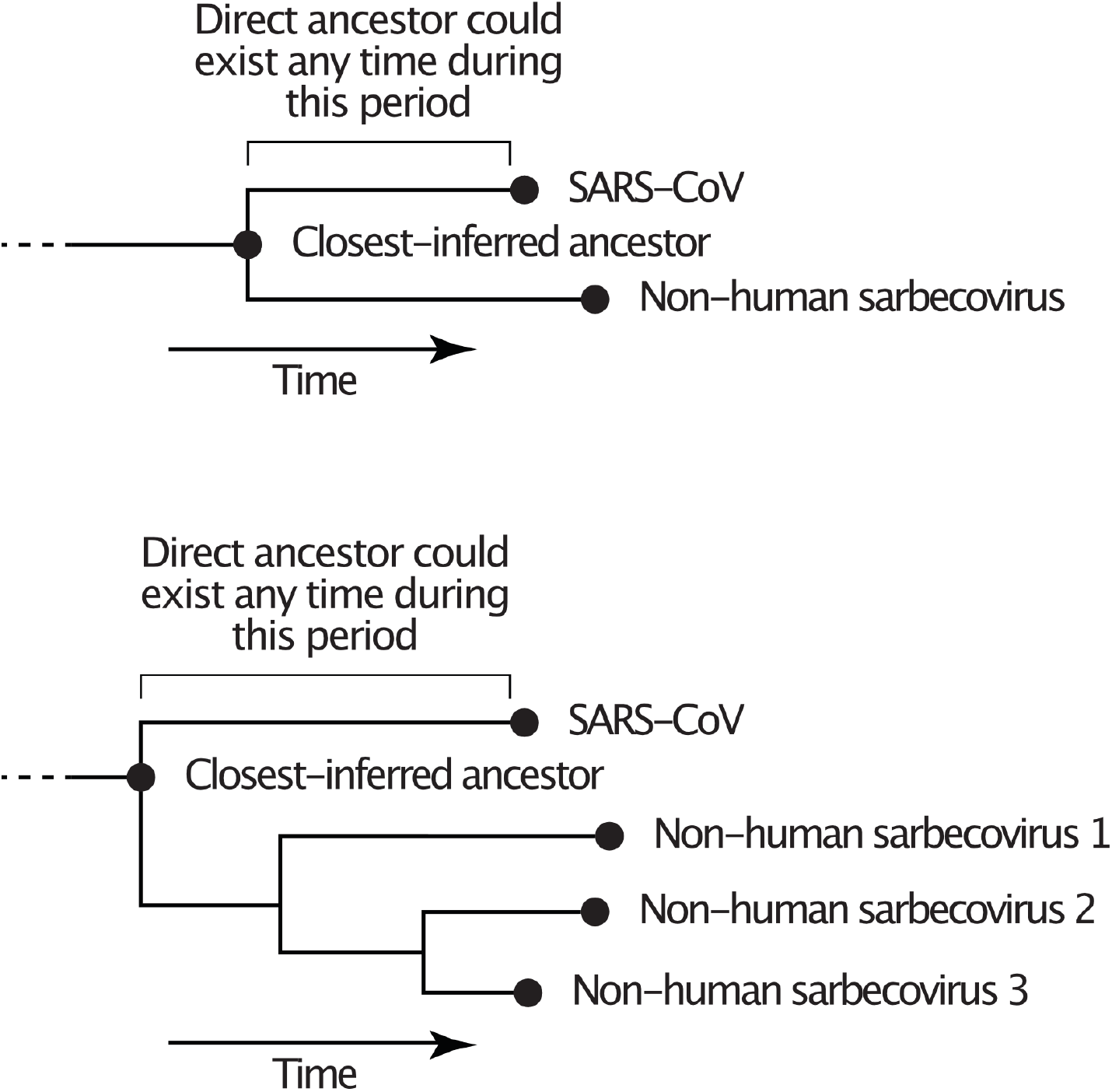
Schematics illustrating the relationship between a SARS-CoV, the direct ancestor, and the closest-inferred ancestor. The closest-inferred ancestor could be the tMRCA of a given SARS-CoV and one non-human sarbecovirus (top) or the tMRCA of the SARS-CoV and multiple non-human sarbecoviruses (bottom); the closest-inferred ancestor is the parent node of the SARS-CoV. The direct ancestor is the bat sarbecovirus that was involved in each cross-species transmission event leading to the SARS-CoV and could have existed at any point on the branch leading from the closest-inferred ancestor to the SARS-CoV.

**Extended Data Fig. 2.**
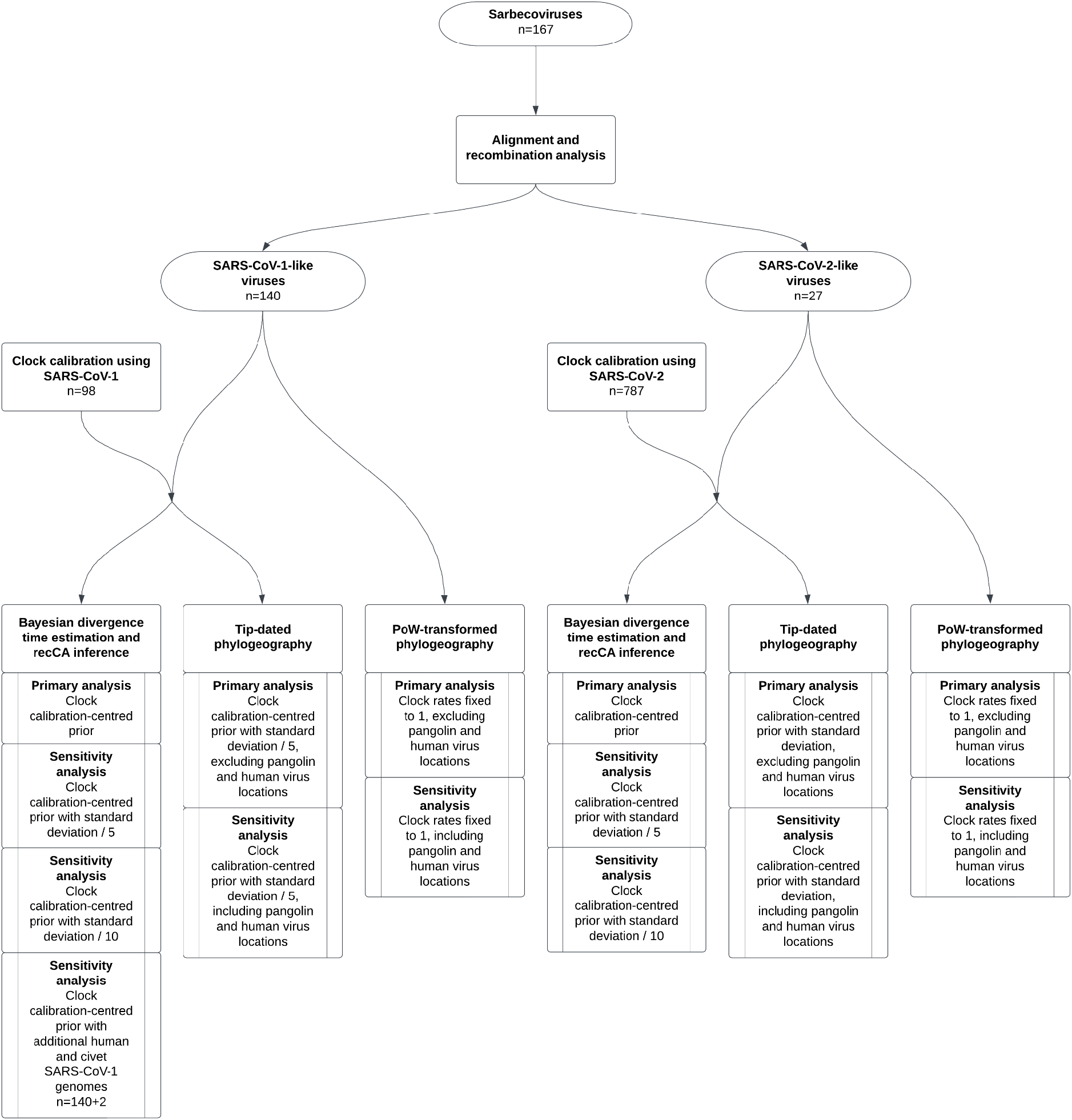
Workflow to perform phylogenetic and phylogeographic analyses. Starting with 167 sarbecovirus genomes, we align and perform recombination analysis to determine the number of SARS-CoV-1-like and SARS-CoV-2-like viruses and their corresponding NRRs for subsequent analyses. We then calibrate the molecular clock for SARS-CoV-1-like viruses and SARS-CoV-2-like viruses using SARS-CoV-1 genomes and SARS-CoV-2 genomes (these datasets are separate from the SARS-CoV-1-like viruses and SARS-CoV-2-like viruses), respectively. Finally, we perform Bayesian divergence time estimation, recCA inference, and phylogeography analyses, each of which comprised both primary and sensitivity analyses. Ovals indicate the number of sarbecovirus genomes included in subsequent steps. Rectangles without inner vertical lines indicate analyses performed; rectangles with inner vertical lines indicate primary versus sensitivity analyses, with non-bolded text describing the details of the analysis performed.

**Extended Data Fig. 3.**
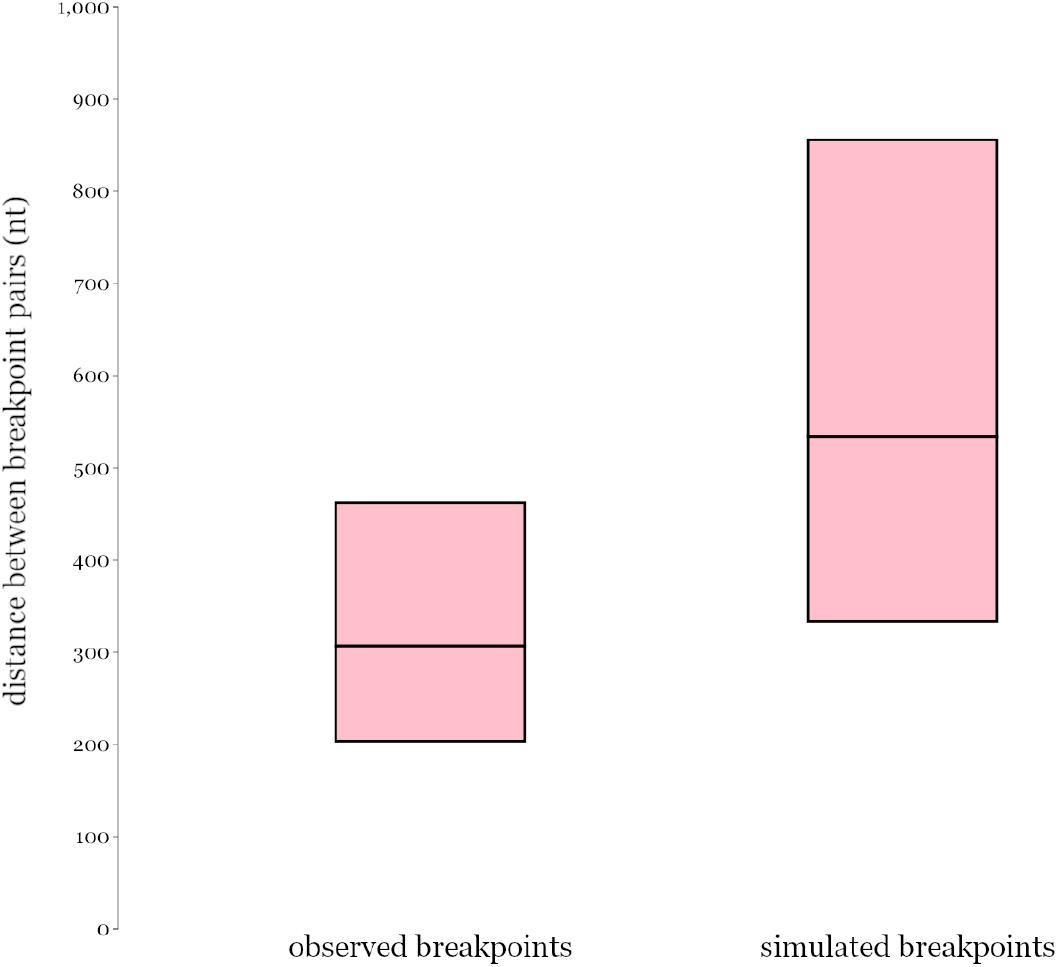
Mean and 90% confidence intervals of the log-normal breakpoint pair distance distributions for the observed SARS-CoV-1-like and SARS-CoV-2-like estimated breakpoints and the simulated randomly sampled breakpoint sets.

**Extended Data Fig. 4.**
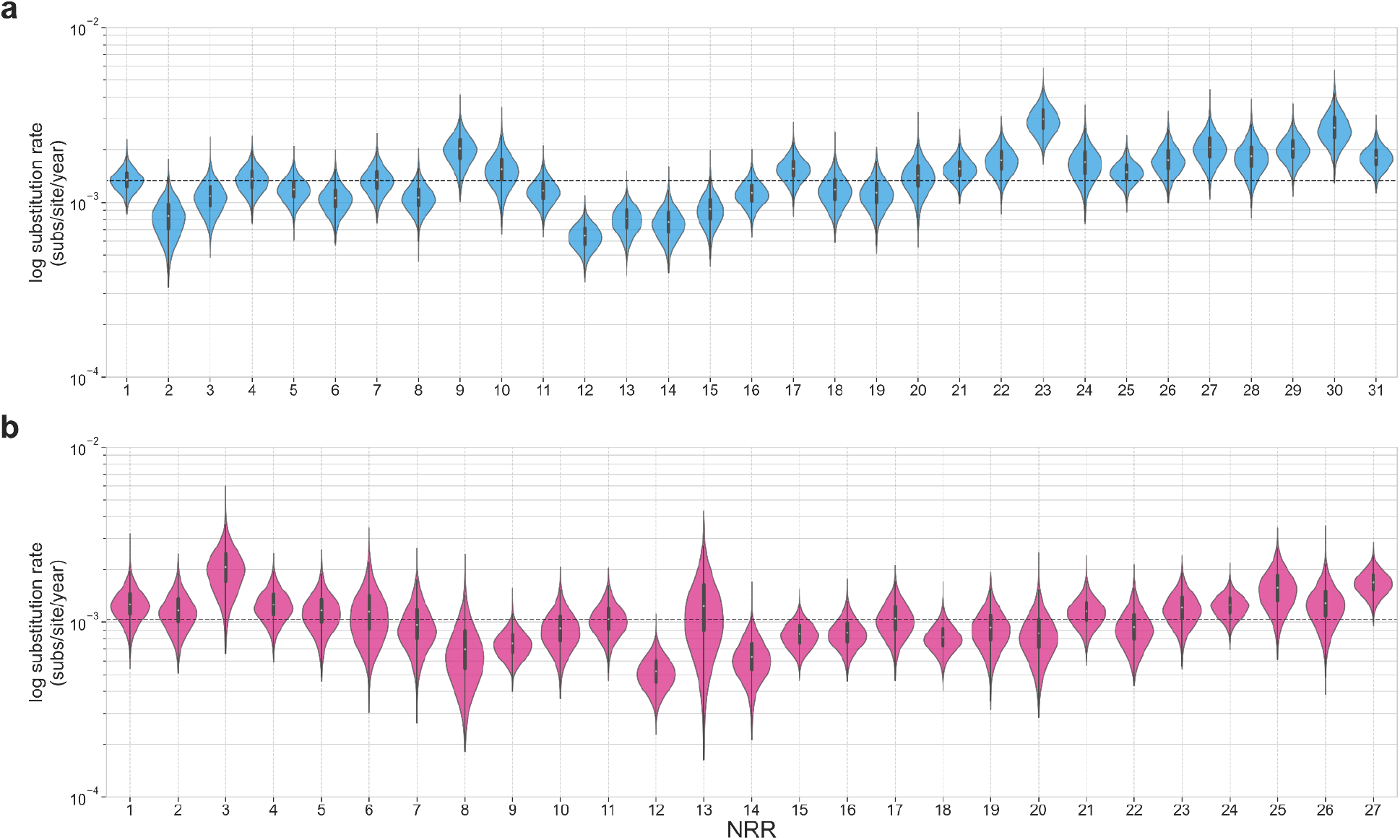
Substitution rates for the SARS-CoV-1-like and SARS-CoV-2-like viruses. Substitution rates (substitutions/site/year) across the (**a**) 31 NRRs for the SARS-CoV-1-like viruses and (**b**) 27 NRRs for the SARS-CoV-2-like viruses. The dashed line in each panel is the respective median substitution rate. The dots and thick lines within the violins indicate the median and interquartile range.

**Extended Data Fig. 5.**
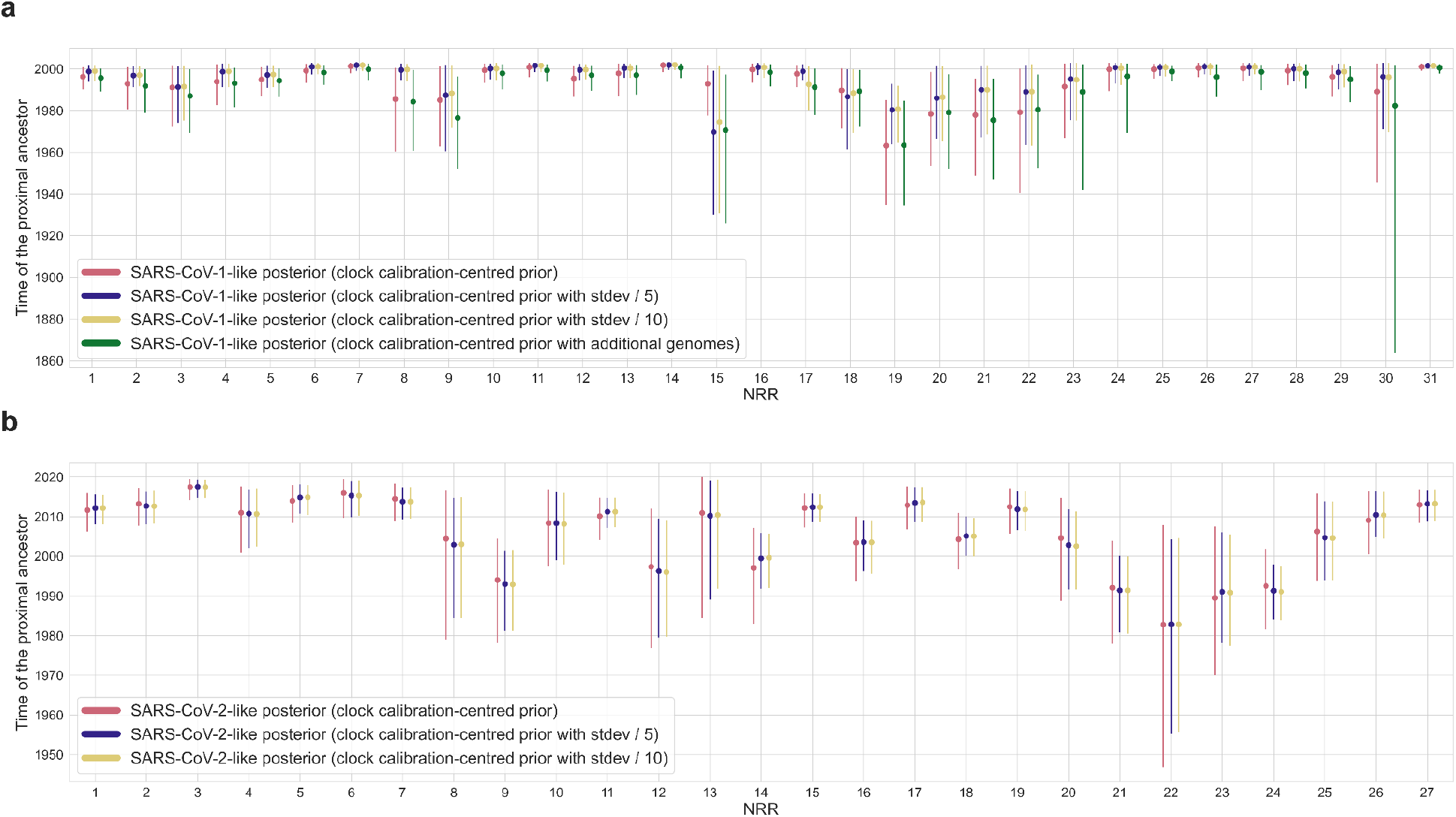
Time of the closest-inferred ancestor sensitivity analysis. **a,** Time of the closest-inferred ancestor of SARS-CoV-1 across the 31 NRRs of the SARS-CoV-1-like viruses with the calibrated molecular clock from the SARS-CoV-1 analysis, the calibrated molecular clock but the standard deviation of the clock rate divided by 5, the calibrated molecular clock but the standard deviation of the clock rate divided by 10, and when adding in one SARS-CoV-1 genome from a civet and an additional SARS-CoV-1 genome from a human. **b,** Time of the closest-inferred ancestor of SARS-CoV-2 across the 27 NRRs of the SARS-CoV-2-like viruses with the calibrated molecular clock from the SARS-CoV-2 analysis, the calibrated molecular clock but the standard deviation of the clock rate divided by 5, and the calibrated molecular clock but the standard deviation of the clock rate divided by 10. The dots indicate the median and the lines are the 95% HPD. Refer to Supplementary Tables 3 and 4 for numerical values.

**Extended Data Fig. 6.**
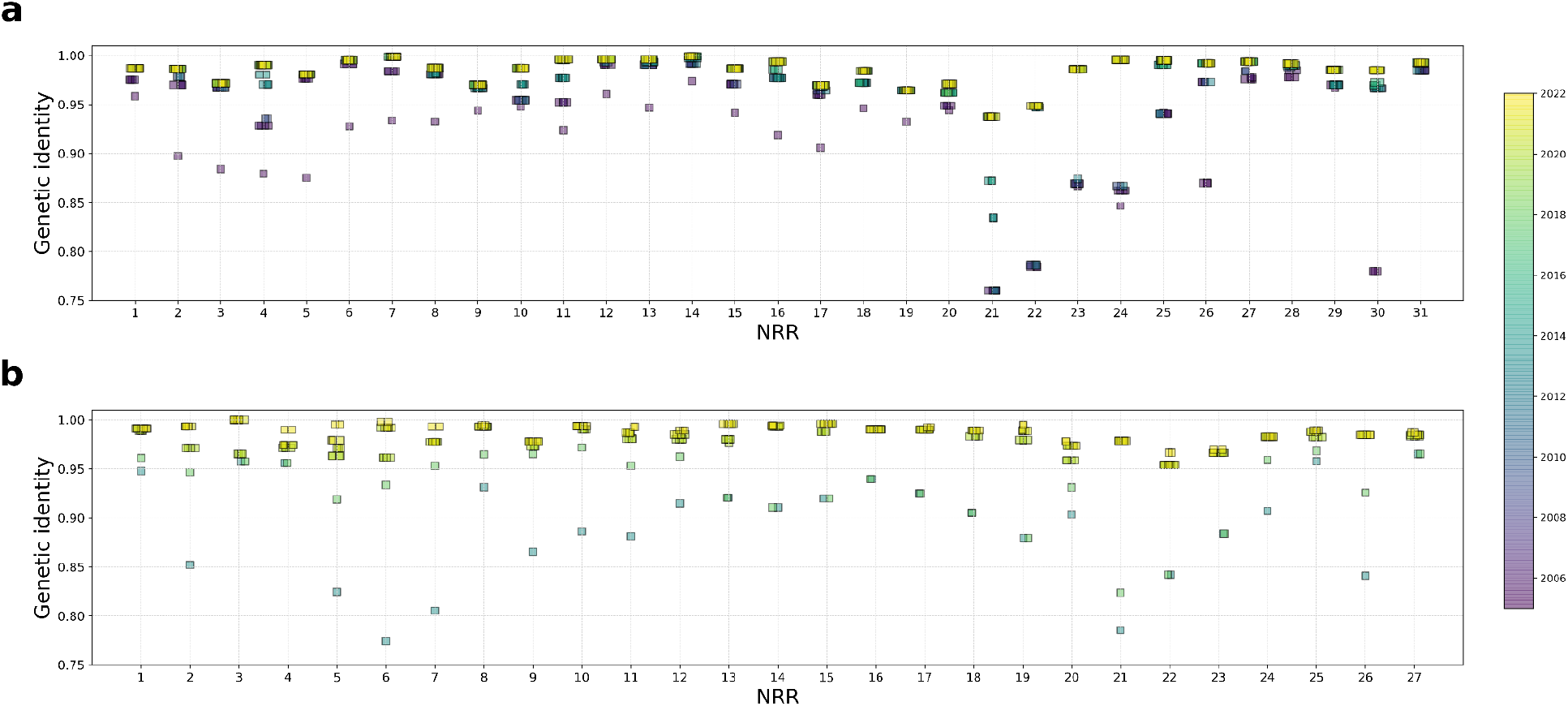
Genetic similarity of the NRRs of the recCA to each SARS-CoV over time. **a,** The similarity of each NRR of the recCA of SARS-CoV-1 to SARS-CoV-1 as a function of time. **b,** The similarity of each NRR of the recCA of SARS-CoV-2 to SARS-CoV-2 as a function of time. Each square represents the similarity of the NRR of the recCA to its relative SARS-CoV at one point in time and is colored by date of the most recent non-human sarbecovirus in the reconstruction. See Fig. 2 for the similarity of the complete recCA of each SARS-CoV to its relative SARS-CoV over time.

**Extended Data Fig. 7.**
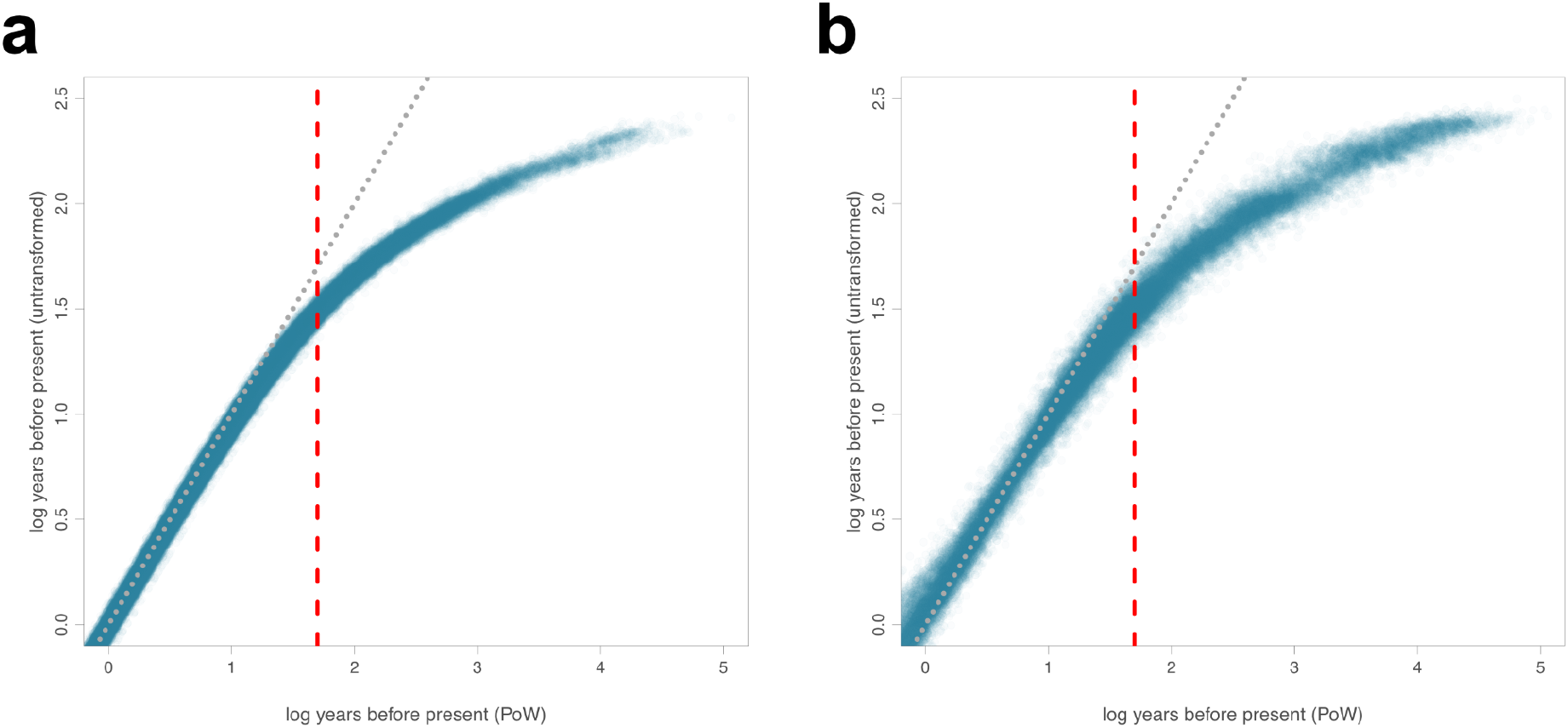
Prisoner of War (PoW) transformation comparison. The log number of years of each internal node before the most recently sampled taxon compared to the log number of years of the same node after applying a PoW transformation for (**a**) 100 SARS-CoV-1-like posterior phylogenies for each NRR and (**b**) 100 SARS-CoV-2-like posterior phylogenies for each NRR. Red dashed line at 50 years before the most recently sampled taxon indicates a threshold beyond which saturation effects substantially influence tMRCA estimates. Diagonal dotted line depicts a line of equality, with a one-to-one correspondence between the two inferences on the two axes.

**Extended Data Fig. 8.**
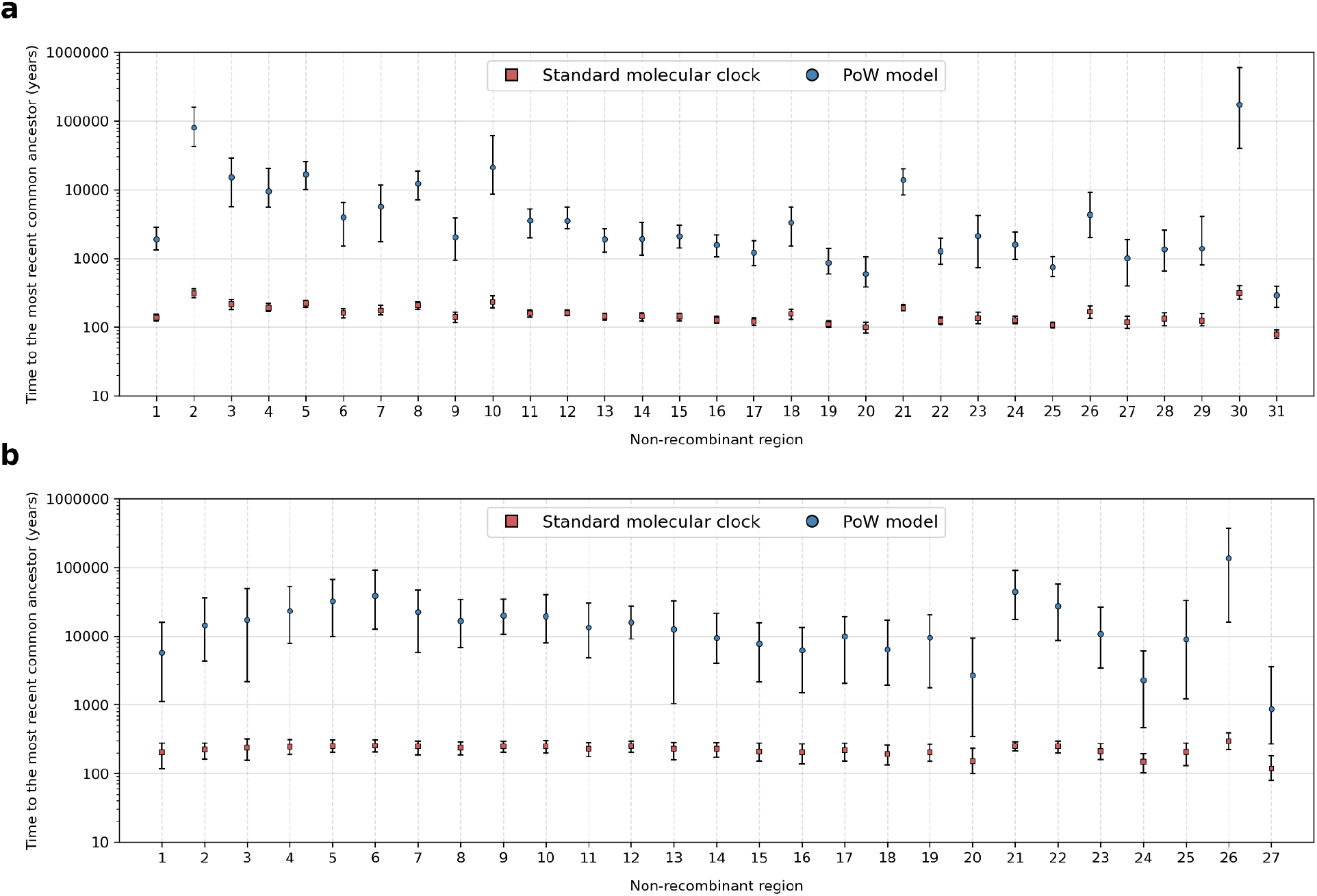
Time to the most recent common ancestor (tMRCA) after PoW transformation. Estimates of time to the most recent common ancestor for the (**a**) SARS-CoV-1-like viruses and (**b**) SARS-CoV-2-like viruses for each NRR when using a standard molecular clock or the PoW model. Each square indicates the median tMRCA and the lines indicate the 95% HPD.

**Extended Data Fig. 9.**
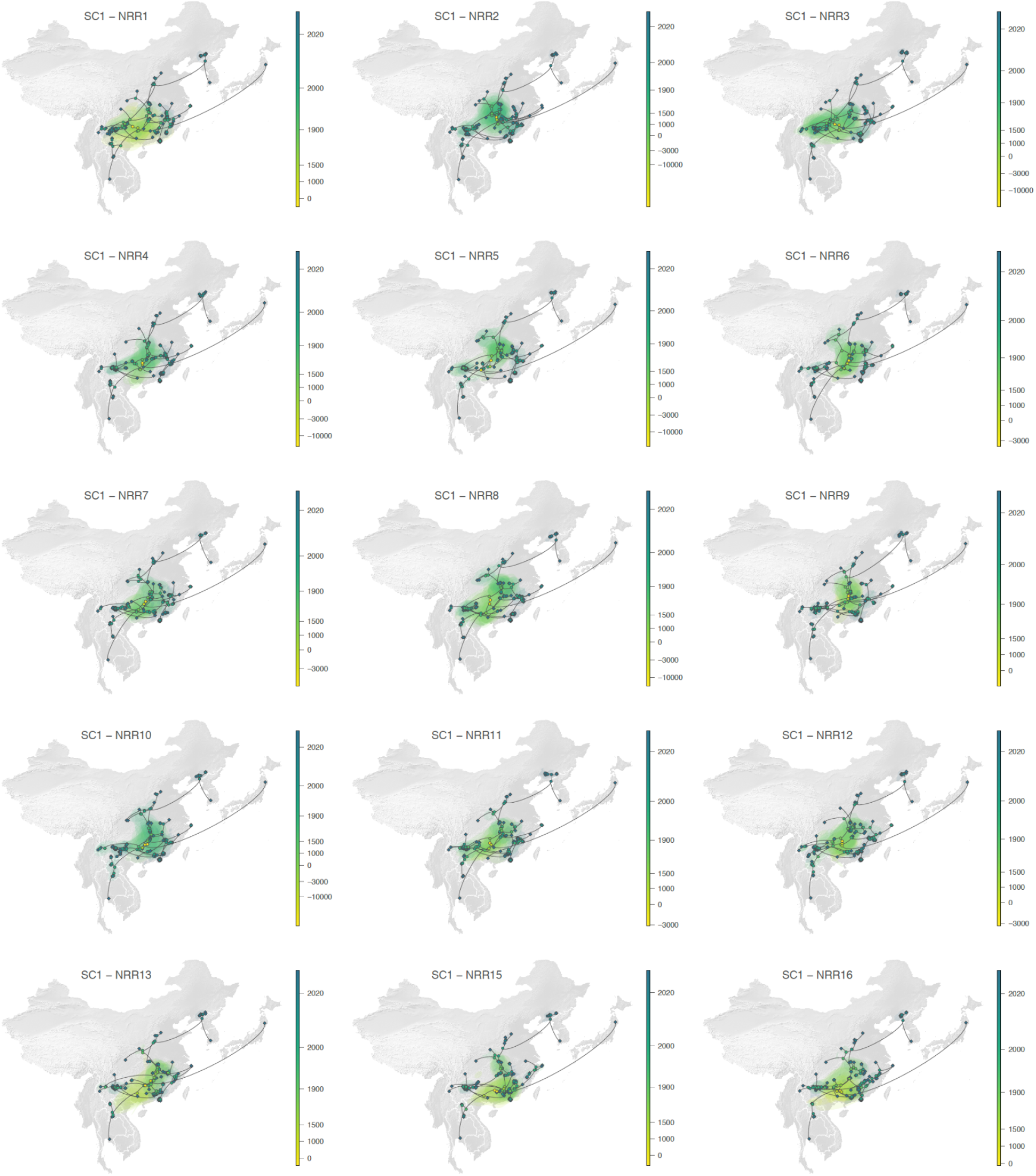
Continuous phylogeographic reconstruction of the dispersal history of viral lineages for the SARS-CoV-1-like (SC1) viruses. We report the reconstructions obtained for NRRs1-13 and 15-16 (NRR14 being reported in the main Fig. 3g; see Extended Data Fig. 10 for NRRs17-31). For each NRR, we map the maximum clade credibility (MCC) tree whose nodes are coloured from yellow (the time of the most recent common ancestor) to blue (most recent collection date) and are superimposed on 80% HPD polygons coloured according to the same log-transformed time scale and reflecting the uncertainty of the Bayesian phylogeographic inference. HPD polygons were only computed and reported for the last 1,000 years. As in Fig. 3g-h, MCC tree tip nodes are displayed as squares (as opposed to dots for internal ones) and dispersal direction (anti-clockwise) of viral lineages is indicated by the edge curvature.

**Extended Data Fig. 10.**
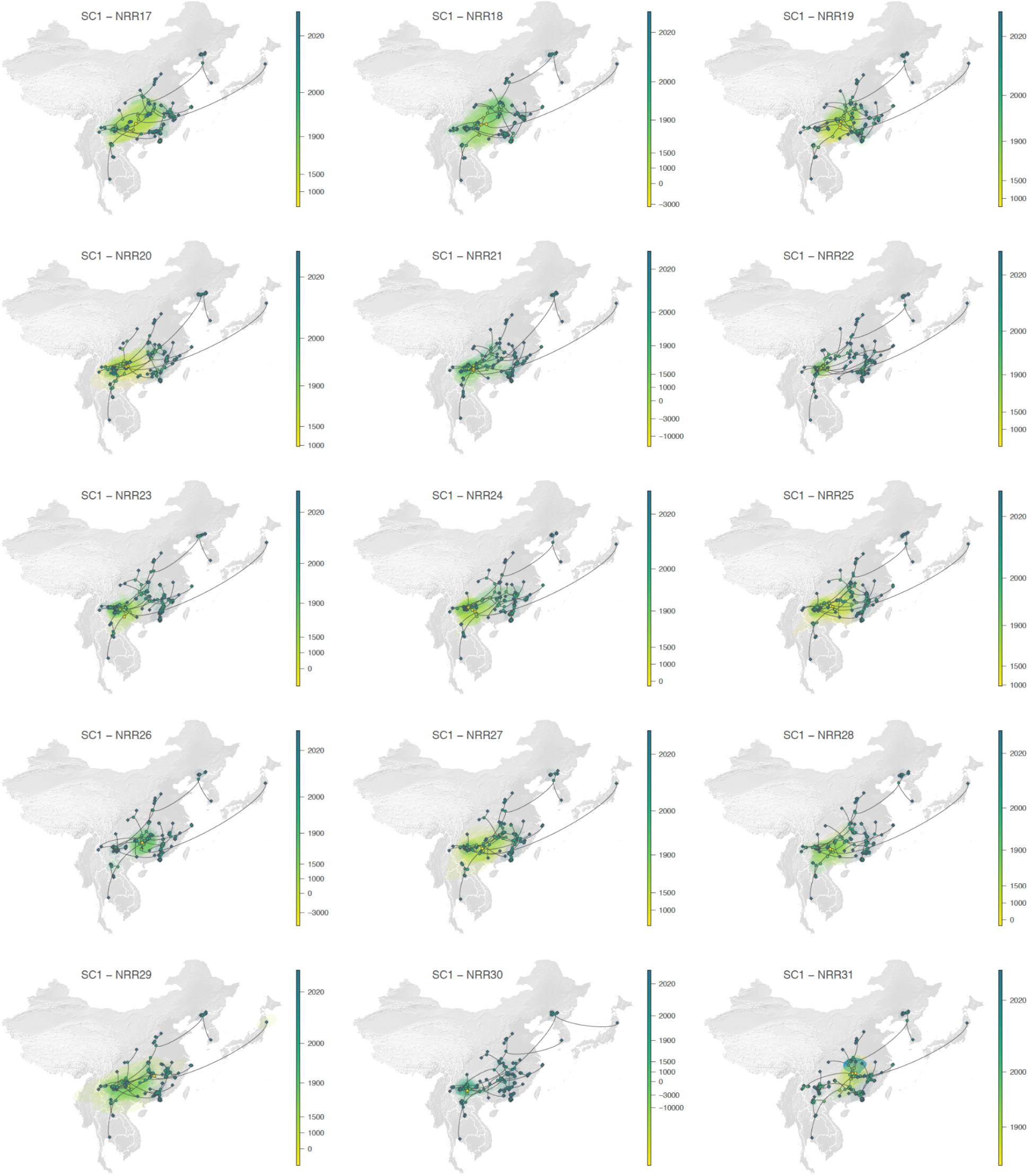
Continuous phylogeographic reconstruction of the dispersal history of viral lineages for the SARS-CoV-1-like (SC1) viruses. We report the reconstructions obtained for NRRs17-31 (see Extended Data Fig. 9 for NRRs 1-13 and 15-17, NRR14 being reported in the main Fig. 3g). For each NRR, we map the maximum clade credibility (MCC) tree whose nodes are coloured from yellow (the time of the most recent common ancestor) to blue (most recent collection date) and are superimposed on 80% HPD polygons coloured according to the same log-transformed time scale and reflecting the uncertainty of the Bayesian phylogeographic inference. HPD polygons were only computed and reported for the last 1,000 years. As in Fig. 3g-h, MCC tree tip nodes are displayed as squares (as opposed to dots for internal ones) and dispersal direction (anti-clockwise) of viral lineages is indicated by the edge curvature.

**Extended Data Fig. 11.**
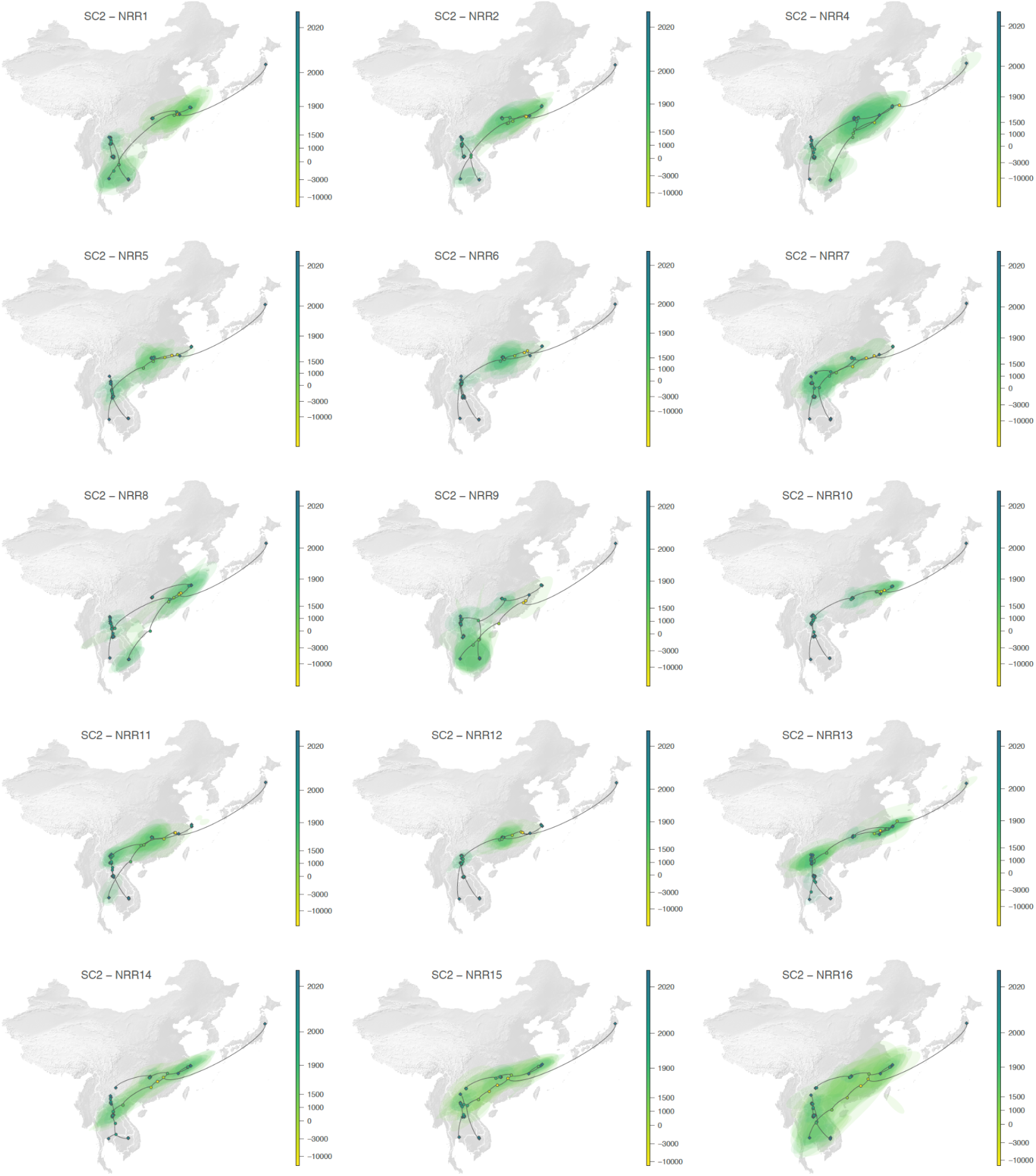
Continuous phylogeographic reconstruction of the dispersal history of viral lineages for the SARS-CoV-2-like (SC2) viruses. We report the reconstructions obtained for NRRs1-2 and 4-16 (NRR3 being reported in the main Fig. 3h; see Extended Data Fig. 12 for NRRs 17-27). For each NRR, we map the maximum clade credibility (MCC) tree whose nodes are coloured from yellow (the time of the most recent common ancestor) to blue (most recent collection date) and are superimposed on 80% HPD polygons coloured according to the same log-transformed time scale and reflecting the uncertainty of the Bayesian phylogeographic inference. HPD polygons were only computed and reported for the last 1,000 years. As in Fig. 3g-h, MCC tree tip nodes are displayed as squares (as opposed to dots for internal ones) and dispersal direction (anti-clockwise) of viral lineages is indicated by the edge curvature.

**Extended Data Fig. 12.**
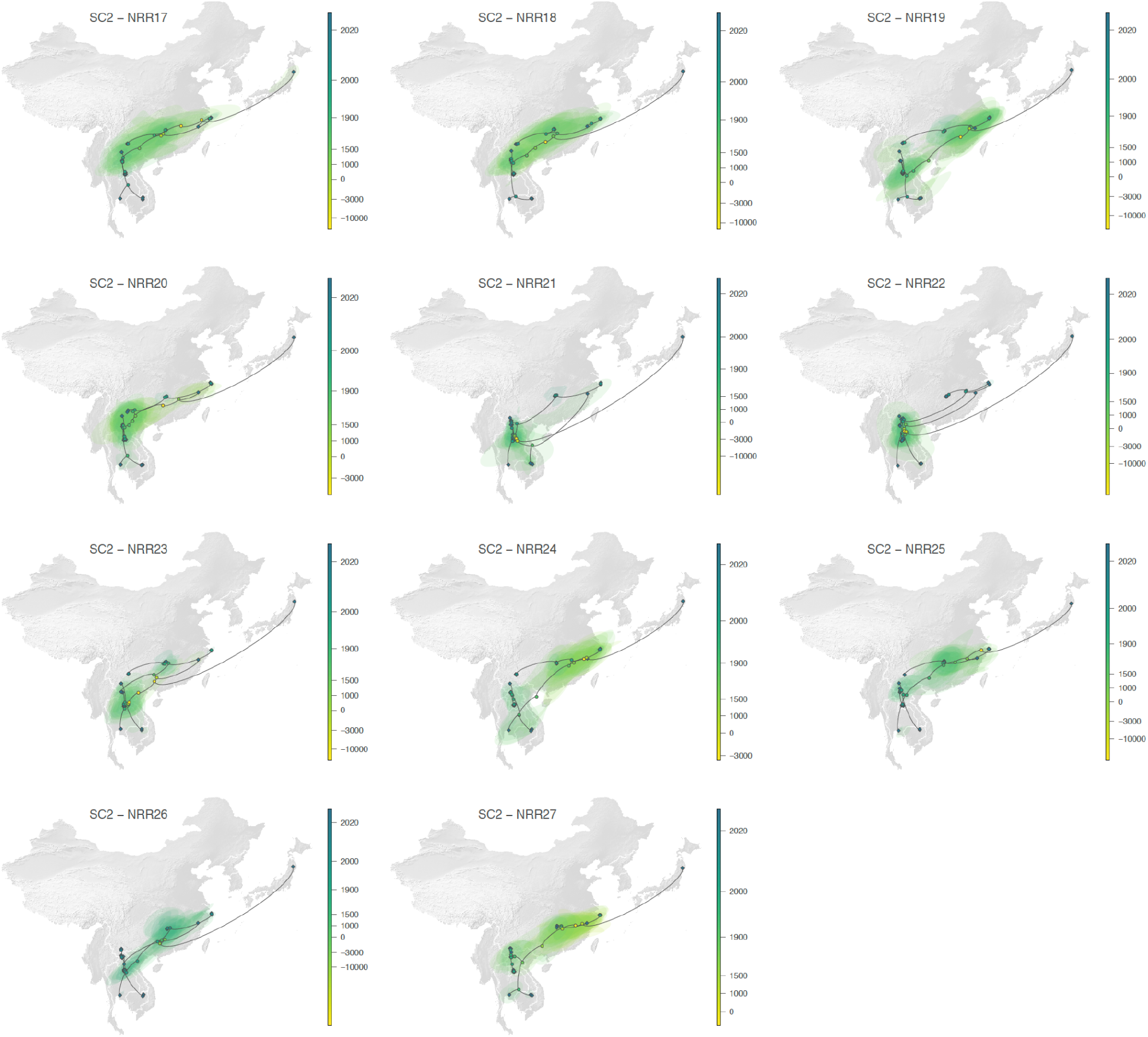
Continuous phylogeographic reconstruction of the dispersal history of viral lineages for the SARS-CoV-2-like (SC2) viruses. We report the reconstructions obtained for NRRs17-27 (see Extended Data Fig. 11 for NRRs 1-2 and 4-16, NRR3 being reported in the main Fig. 3h). For each NRR, we map the maximum clade credibility (MCC) tree whose nodes are coloured from yellow (the time of the most recent common ancestor) to blue (most recent collection date) and are superimposed on 80% HPD polygons coloured according to the same log-transformed time scale and reflecting the uncertainty of the Bayesian phylogeographic inference. HPD polygons were only computed and reported for the last 1,000 years. As in Fig. 3g-h, MCC tree tip nodes are displayed as squares (as opposed to dots for internal ones) and dispersal direction (anti-clockwise) of viral lineages is indicated by the edge curvature.

**Extended Data Fig. 13.**
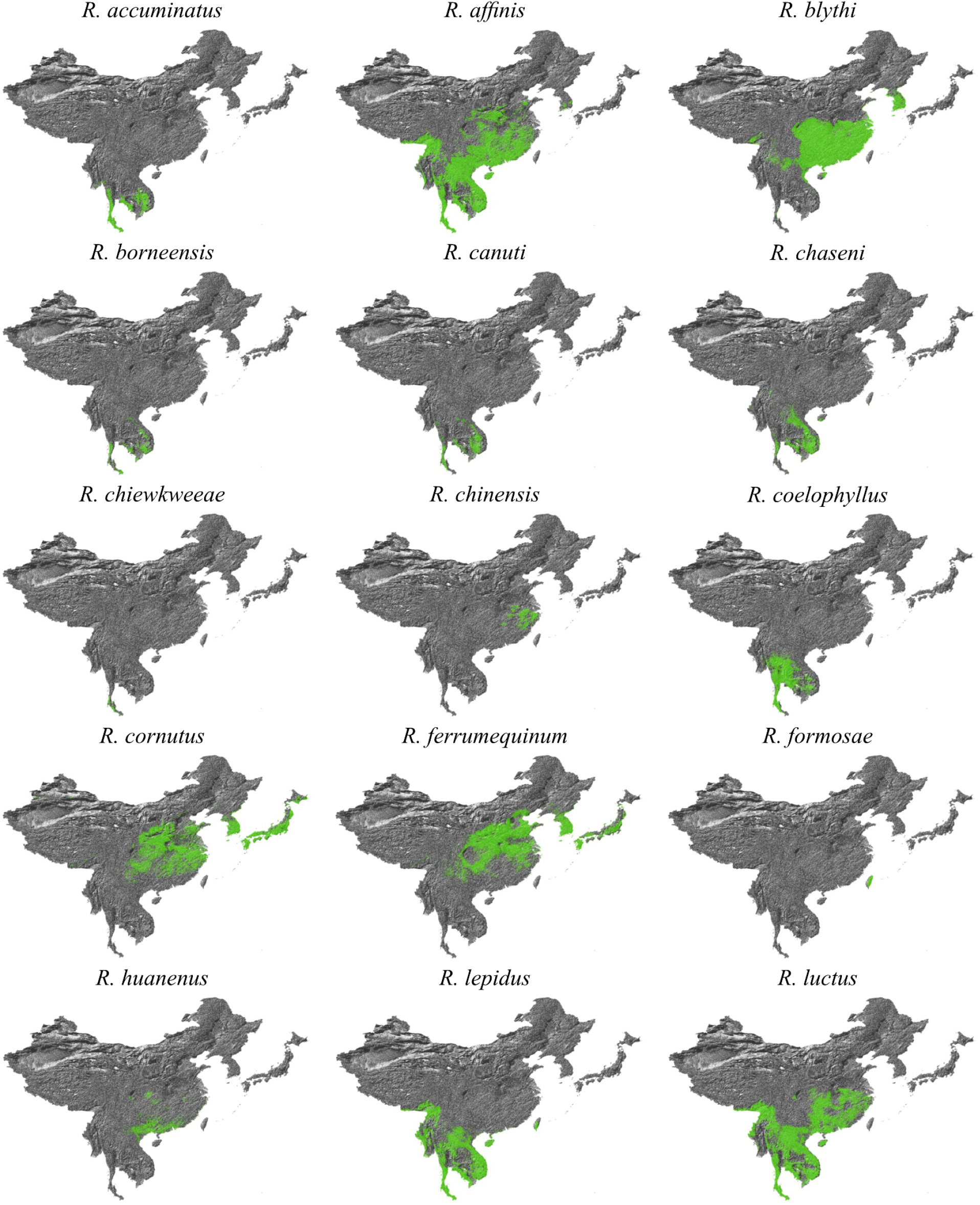
Distribution maps for species of the *Rhinolophus* genus. See Extended Data Fig. 14–15 for remaining bat distribution maps.

**Extended Data Fig. 14.**
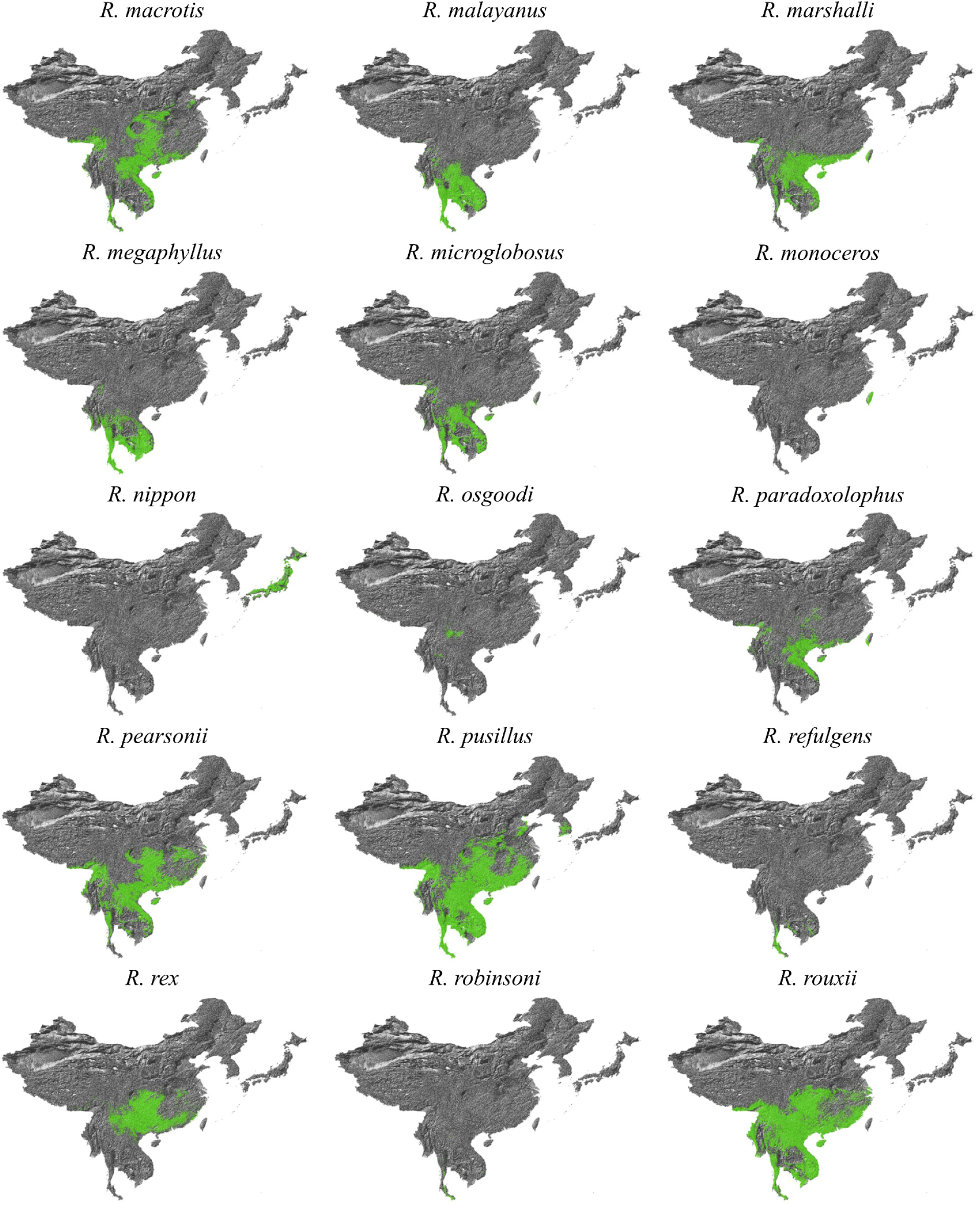
Distribution maps for species of the *Rhinolophus* genus. See Extended Data Fig. 13 and 15 for remaining bat distribution maps.

**Extended Data Fig. 15.**
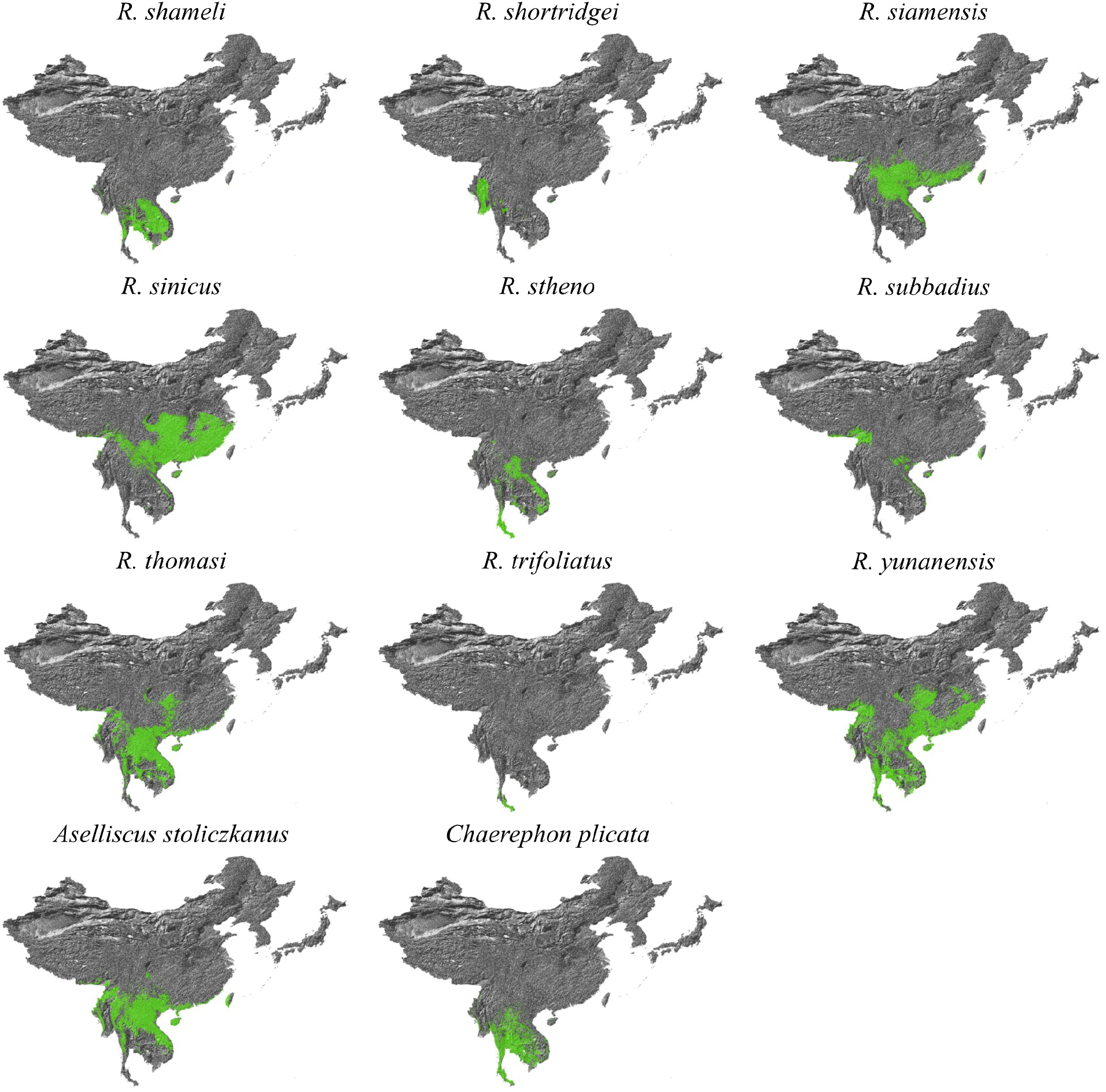
Distribution maps for species of the *Rhinolophus*, *Ascelliscus*, and *Chaerephon* genera. Species from the *Ascelliscus* and *Chaerephon* genera are limited to those that were hosts to sarbecoviruses in our analyses. See Extended Data Fig. 13–14 for remaining bat distribution maps.

**Extended Data Fig. 16.**
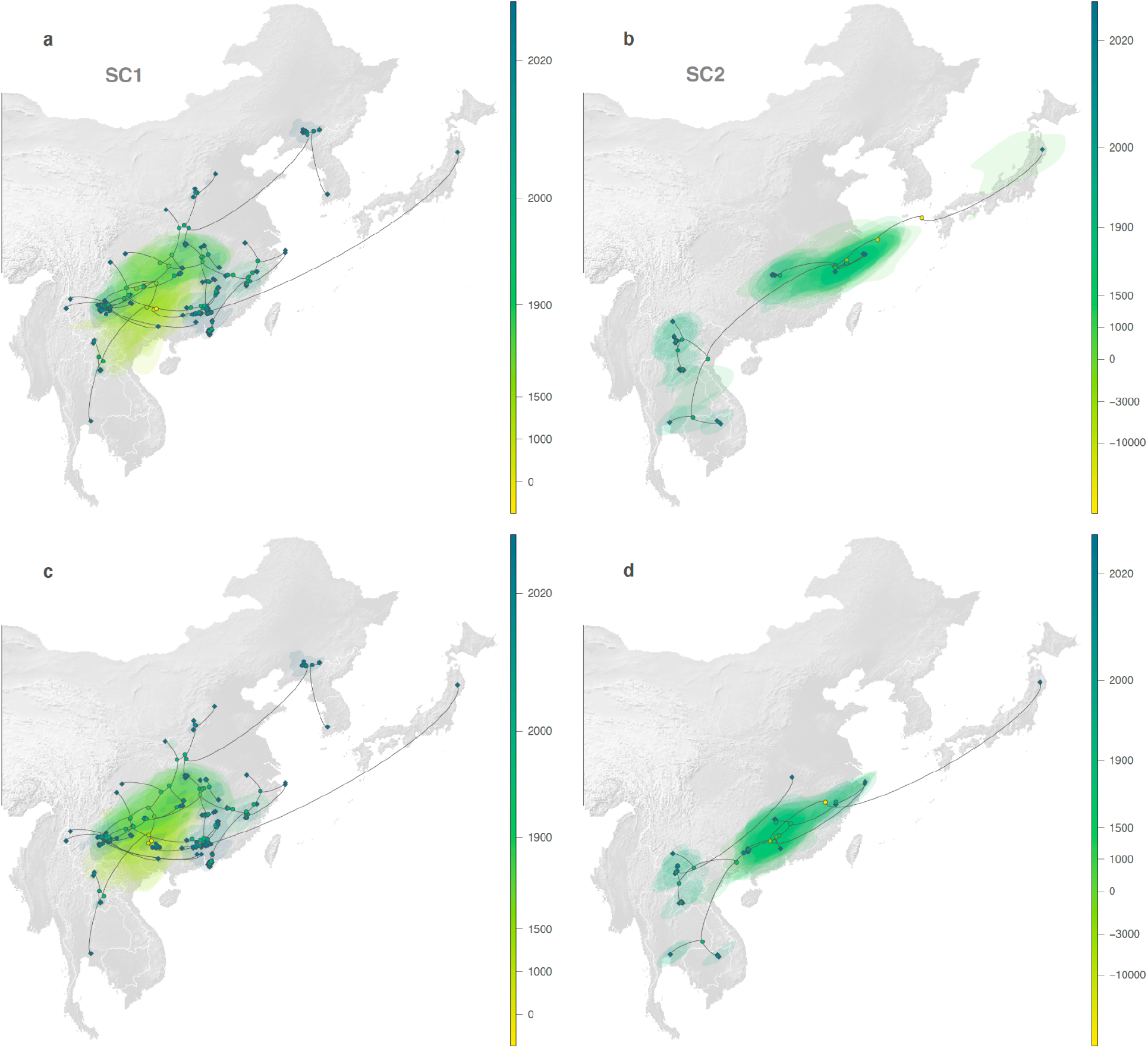
Comparison of the continuous phylogeographic reconstructions for two selected NRRSs (NRR14 for SARS-CoV-1-like viruses [SC1] on the left, and NRR3 for SARS-CoV-2-like viruses [SC2] on the right) when not considering (a-b) or when considering (c-d) the human and pangolin sequences in the analyses. For each reconstruction, we map the corresponding MCC tree whose nodes are coloured from yellow (the time of the most recent common ancestor) to blue (most recent collection date) and are superimposed on 80% HPD polygons coloured according to a log-transformed time scale and reflecting the uncertainty of the Bayesian phylogeographic inference. HPD polygons were only computed and reported for the last 1,000 years. As in Fig. 3g-h, MCC tree tip nodes are displayed as squares (as opposed to dots for internal ones) and dispersal direction (anti-clockwise) of viral lineages is indicated by the edge curvature.

**Extended Data Fig. 17.**
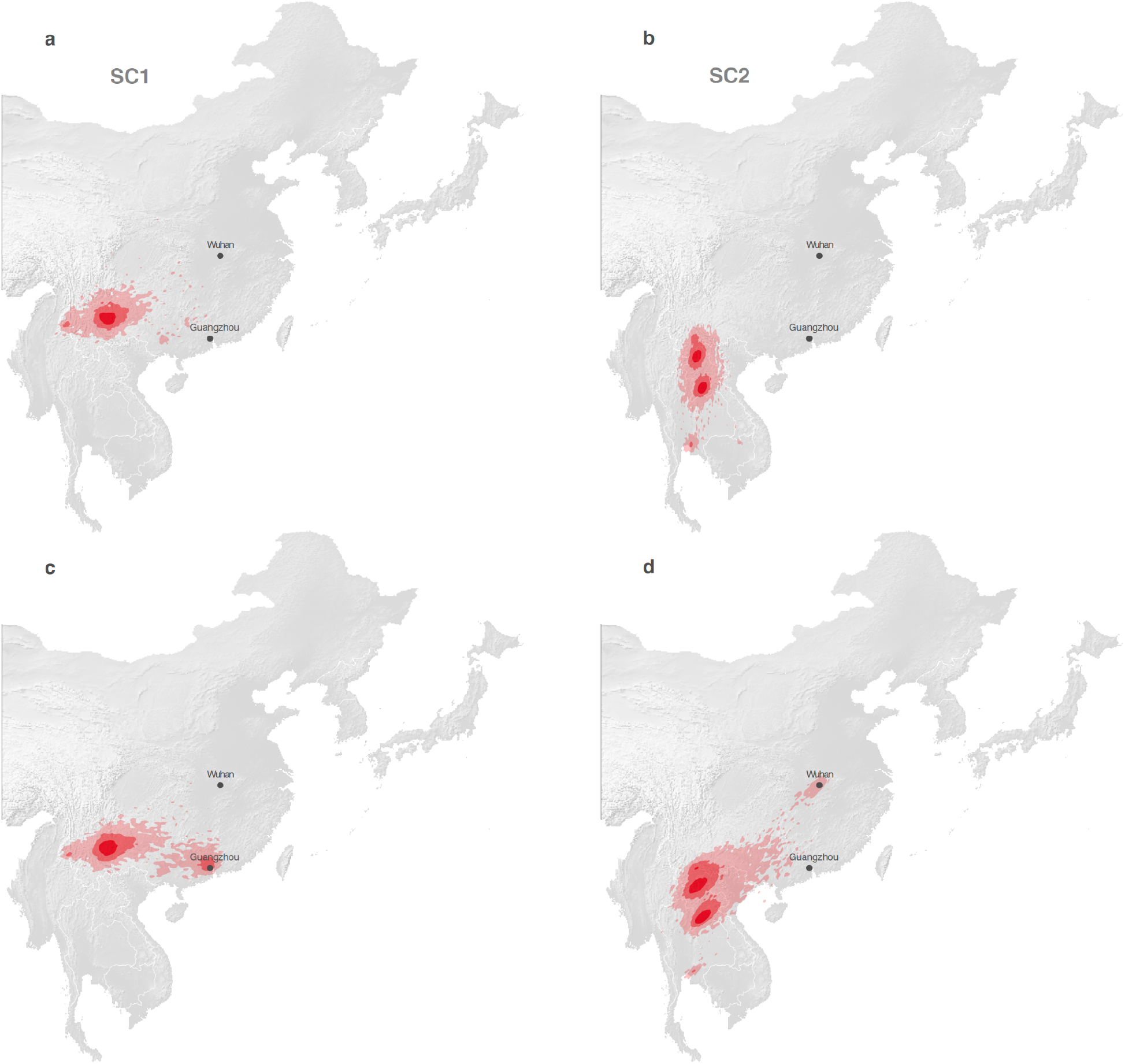
Comparison of the estimated position of the recCA according to continuous phylogeographic reconstructions that did not consider (a-b) or did consider (c-d) the human and pangolin sequences in the analyses. As in Figure 4g-h, the estimated position of the recCA is based on the continuous phylogeographic reconstruction of all NRRs. We here display the 95%, 75% and 50% HPD regions with an increasing red color darkness. “G.” and “W.” indicate the position of the cities of Guangzhou and Wuhan, respectively.

**Extended Data Fig. 18.**
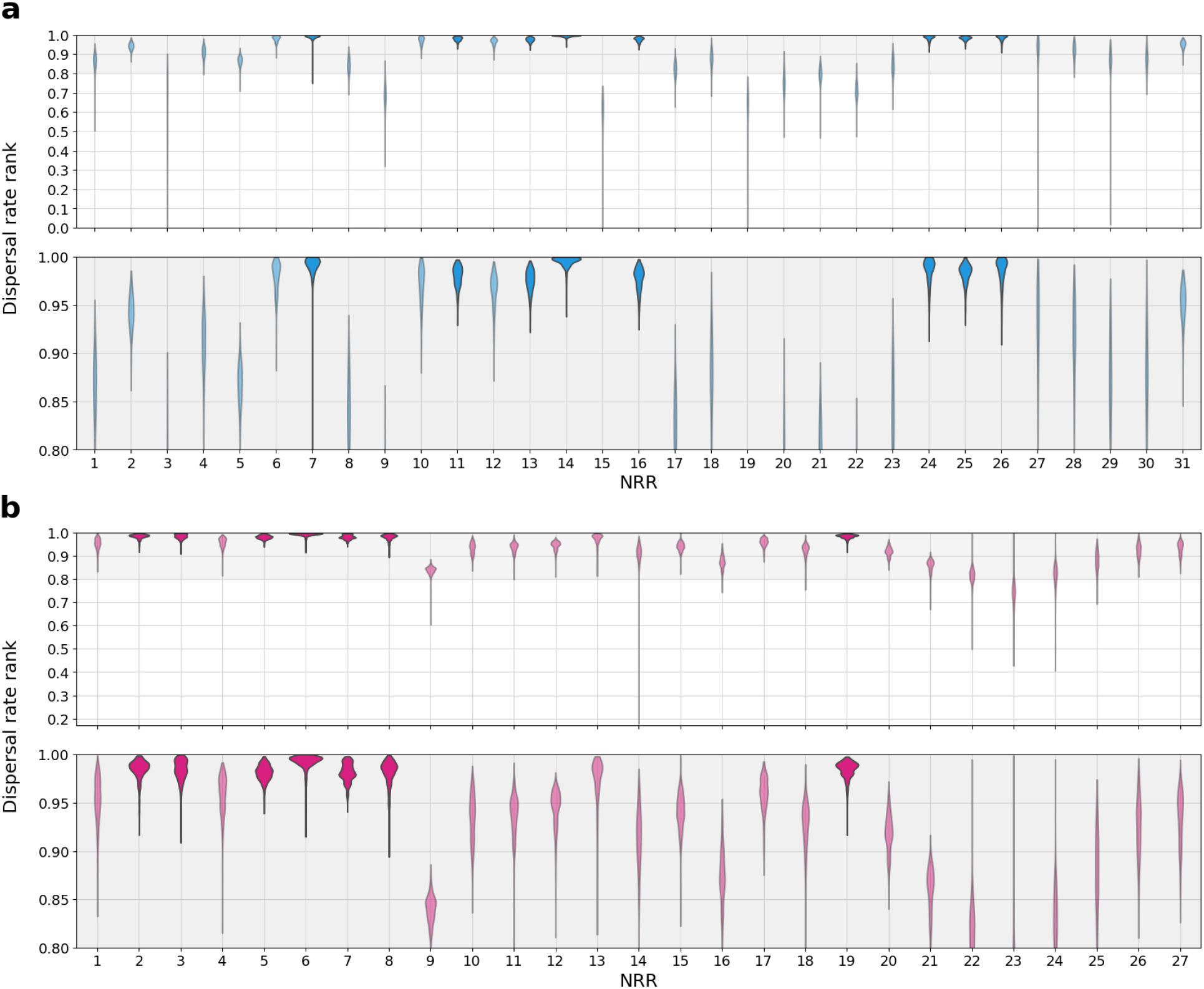
Ranking of dispersal velocity for the closest-inferred bat virus ancestor to reach the province of emergence. **a,** The rank for the dispersal velocity necessary to travel from the location of the closest-inferred bat virus ancestor of SARS-CoV-1 from each NRR to the nearest point in Guangdong Province, given the time between emergence and this ancestor and the bat virus dispersal velocities for each NRR. **b,** The rank for the dispersal velocity necessary to travel from the location of the closest-inferred bat virus ancestor of SARS-CoV-2 from each NRR to the nearest point in Hubei Province, given the time between emergence and this ancestor and the bat virus dispersal velocities for each NRR. The top 20% of the dispersal rate ranks are zoomed-in upon in the lower panels of **a** and **b**. Violins are darkened if at least 95% of the dispersal velocity rank posterior is greater than 0.95.

## Supplementary Information

**Supplementary Data File 1: Sarbecovirus genomes metadata.** Temporal, geographic and acknowledgement information for the 167 sarbecovirus genomes used in the paper. Sequences used for each SARS-CoV-1-like and SARS-CoV-2-like GARD recombination analysis and the virus group of each genome (SARS-CoV-1-like, SARS-CoV-2-like or outgroup) are also indicated in the table.

**Supplementary Data File 2: Genome accessions.** Genome accessions for all sarbecovirus genomes used in the paper.

**Supplementary Table 1.** Posterior clock rates across each NRR from the clock calibration using SARS-CoV-1 genomes and Bayesian divergence time analyses of SARS-CoV-1-like viruses. The clock calibration posterior results are used as a prior for the Bayesian divergence time analyses, with the standard deviation of the clock calibration results varying across the analyses.

| NRR | Clock calibration<sup>a</sup> | Bayesian divergence time estimation clock posterior<sup>b</sup> |  |  |
| --- | --- | --- | --- | --- |
|  |  | Primary analysis | Robustness analysis |  |
|  |  | Clock calibration-centred prior | Clock calibration-centred prior with stdev / 5 | Clock calibration-centred prior with stdev / 10 |
| 1 | 2.06e-03 (3.81e-04) | 1.35e-03<br>(1.01e-03, 1.72e-03) | 2.00e-03<br>(1.86e-03, 2.14e-03) | 2.04e-03<br>(1.97e-03, 2.12e-03) |
| 2 | 1.21e-03 (3.74e-04) | 8.36e-04<br>(4.88e-04, 1.23e-03) | 1.18e-03<br>(1.04e-03, 1.31e-03) | 1.20e-03<br>(1.13e-03, 1.27e-03) |
| 3 | 1.22e-03 (5.34e-04) | 1.09e-03<br>(6.95e-04, 1.54e-03) | 1.20e-03<br>(1.02e-03, 1.40e-03) | 1.22e-03<br>(1.12e-03, 1.32e-03) |
| 4 | 2.55e-03 (6.45e-04) | 1.36e-03<br>(9.69e-04, 1.84e-03) | 2.41e-03<br>(2.18e-03, 2.65e-03) | 2.51e-03<br>(2.38e-03, 2.64e-03) |
| 5 | 1.61e-03 (3.77e-04) | 1.19e-03<br>(8.63e-04, 1.57e-03) | 1.57e-03<br>(1.43e-03, 1.70e-03) | 1.60e-03<br>(1.53e-03, 1.67e-03) |
| 6 | 2.04e-03 (5.27e-04) | 1.06e-03<br>(7.43e-04, 1.45e-03) | 1.93e-03<br>(1.74e-03, 2.13e-03) | 2.01e-03<br>(1.91e-03, 2.11e-03) |
| 7 | 2.23e-03 (6.06e-04) | 1.35e-03<br>(9.29e-04, 1.80e-03) | 2.12e-03<br>(1.90e-03, 2.34e-03) | 2.20e-03<br>(2.09e-03, 2.32e-03) |
| 8 | 1.85e-03 (4.05e-04) | 1.07e-03<br>(7.53e-04, 1.43e-03) | 1.79e-03<br>(1.65e-03, 1.95e-03) | 1.84e-03<br>(1.76e-03, 1.91e-03) |
| 9 | 2.53e-03 (6.94e-04) | 2.03e-03<br>(1.29e-03, 2.77e-03) | 2.46e-03<br>(2.21e-03, 2.74e-03) | 2.51e-03<br>(2.38e-03, 2.65e-03) |
| 10 | 2.09e-03 (8.46e-04) | 1.54e-03<br>(9.50e-04, 2.22e-03) | 2.01e-03<br>(1.71e-03, 2.31e-03) | 2.06e-03<br>(1.91e-03, 2.23e-03) |
| 11 | 2.34e-03 (4.53e-04) | 1.16e-03<br>(8.38e-04, 1.55e-03) | 2.25e-03<br>(2.09e-03, 2.43e-03) | 2.31e-03<br>(2.23e-03, 2.40e-03) |
| 12 | 1.15e-03 (2.49e-04) | 6.43e-04<br>(4.57e-04, 8.42e-04) | 1.10e-03<br>(1.00e-03, 1.19e-03) | 1.13e-03<br>(1.09e-03, 1.18e-03) |
| 13 | 1.16e-03 (2.50e-04) | 8.09e-04<br>(5.40e-04, 1.09e-03) | 1.13e-03<br>(1.04e-03, 1.23e-03) | 1.15e-03<br>(1.11e-03, 1.20e-03) |
| 14 | 9.28e-04 (2.96e-04) | 7.70e-04<br>(4.91e-04, 1.09e-03) | 9.08e-04<br>(8.04e-04, 1.02e-03) | 9.23e-04<br>(8.66e-04, 9.81e-04) |
| 15 | 1.03e-03 (3.03e-04) | 9.13e-04<br>(5.93e-04, 1.27e-03) | 1.03e-03<br>(9.19e-04, 1.15e-03) | 1.03e-03<br>(9.70e-04, 1.09e-03) |
| 16 | 1.61e-03 (3.32e-04) | 1.13e-03<br>(8.28e-04, 1.50e-03) | 1.57e-03<br>(1.45e-03, 1.70e-03) | 1.60e-03<br>(1.54e-03, 1.66e-03) |
| 17 | 1.87e-03 (4.33e-04) | 1.56e-03<br>(1.13e-03, 2.04e-03) | 1.83e-03<br>(1.68e-03, 2.00e-03) | 1.87e-03<br>(1.78e-03, 1.95e-03) |
| 18 | 1.08e-03 (4.71e-04) | 1.18e-03<br>(7.70e-04, 1.66e-03) | 1.13e-03<br>(9.54e-04, 1.31e-03) | 1.09e-03<br>(1.00e-03, 1.18e-03) |
| 19 | 1.99e-03 (5.10e-04) | 1.13e-03<br>(7.41e-04, 1.59e-03) | 1.91e-03<br>(1.72e-03, 2.09e-03) | 1.97e-03<br>(1.88e-03, 2.07e-03) |
| 20 | 2.21e-03 (7.49e-04) | 1.42e-03<br>(9.18e-04, 2.04e-03) | 2.11e-03<br>(1.85e-03, 2.39e-03) | 2.18e-03<br>(2.04e-03, 2.32e-03) |
| 21 | 3.38e-03 (6.44e-04) | 1.56e-03<br>(1.18e-03, 2.02e-03) | 3.24e-03<br>(3.02e-03, 3.50e-03) | 3.35e-03<br>(3.22e-03, 3.47e-03) |
| 22 | 3.38e-03 (7.33e-04) | 1.74e-03<br>(1.18e-03, 2.34e-03) | 3.24e-03<br>(2.99e-03, 3.54e-03) | 3.34e-03<br>(3.21e-03, 3.49e-03) |
| 23 | 4.00e-03 (1.05e-03) | 2.98e-03<br>(2.05e-03, 4.22e-03) | 3.86e-03<br>(3.47e-03, 4.26e-03) | 3.97e-03<br>(3.77e-03, 4.16e-03) |
| 24 | 2.08e-03 (5.97e-04) | 1.70e-03<br>(1.09e-03, 2.45e-03) | 2.04e-03<br>(1.82e-03, 2.27e-03) | 2.07e-03<br>(1.96e-03, 2.19e-03) |
| 25 | 2.80e-03 (5.11e-04) | 1.49e-03<br>(1.14e-03, 1.94e-03) | 2.70e-03<br>(2.52e-03, 2.90e-03) | 2.78e-03<br>(2.68e-03, 2.87e-03) |
| 26 | 3.15e-03 (9.71e-04) | 1.75e-03<br>(1.21e-03, 2.40e-03) | 2.96e-03<br>(2.62e-03, 3.31e-03) | 3.10e-03<br>(2.92e-03, 3.29e-03) |
| 27 | 3.78e-03 (8.66e-04) | 2.07e-03<br>(1.43e-03, 2.89e-03) | 3.66e-03<br>(3.35e-03, 4.00e-03) | 3.75e-03<br>(3.59e-03, 3.92e-03) |
| 28 | 2.41e-03 (6.81e-04) | 1.83e-03<br>(1.18e-03, 2.59e-03) | 2.36e-03<br>(2.11e-03, 2.61e-03) | 2.40e-03<br>(2.27e-03, 2.53e-03) |
| 29 | 3.22e-03 (8.02e-04) | 2.02e-03<br>(1.42e-03, 2.69e-03) | 3.10e-03<br>(2.81e-03, 3.41e-03) | 3.19e-03<br>(3.04e-03, 3.35e-03) |
| 30 | 6.63e-03 (1.71e-03) | 2.68e-03<br>(1.72e-03, 3.77e-03) | 6.28e-03<br>(5.66e-03, 6.91e-03) | 6.54e-03<br>(6.21e-03, 6.87e-03) |
| 31 | 2.88e-03 (4.98e-04) | 1.81e-03<br>(1.34e-03, 2.34e-03) | 2.81e-03<br>(2.62e-03, 3.00e-03) | 2.87e-03<br>(2.77e-03, 2.96e-03) |

<sup>a</sup>Mean and standard deviation in parentheses
<sup>b</sup>Median and 95% HPD in parentheses

**Supplementary Table 2.**
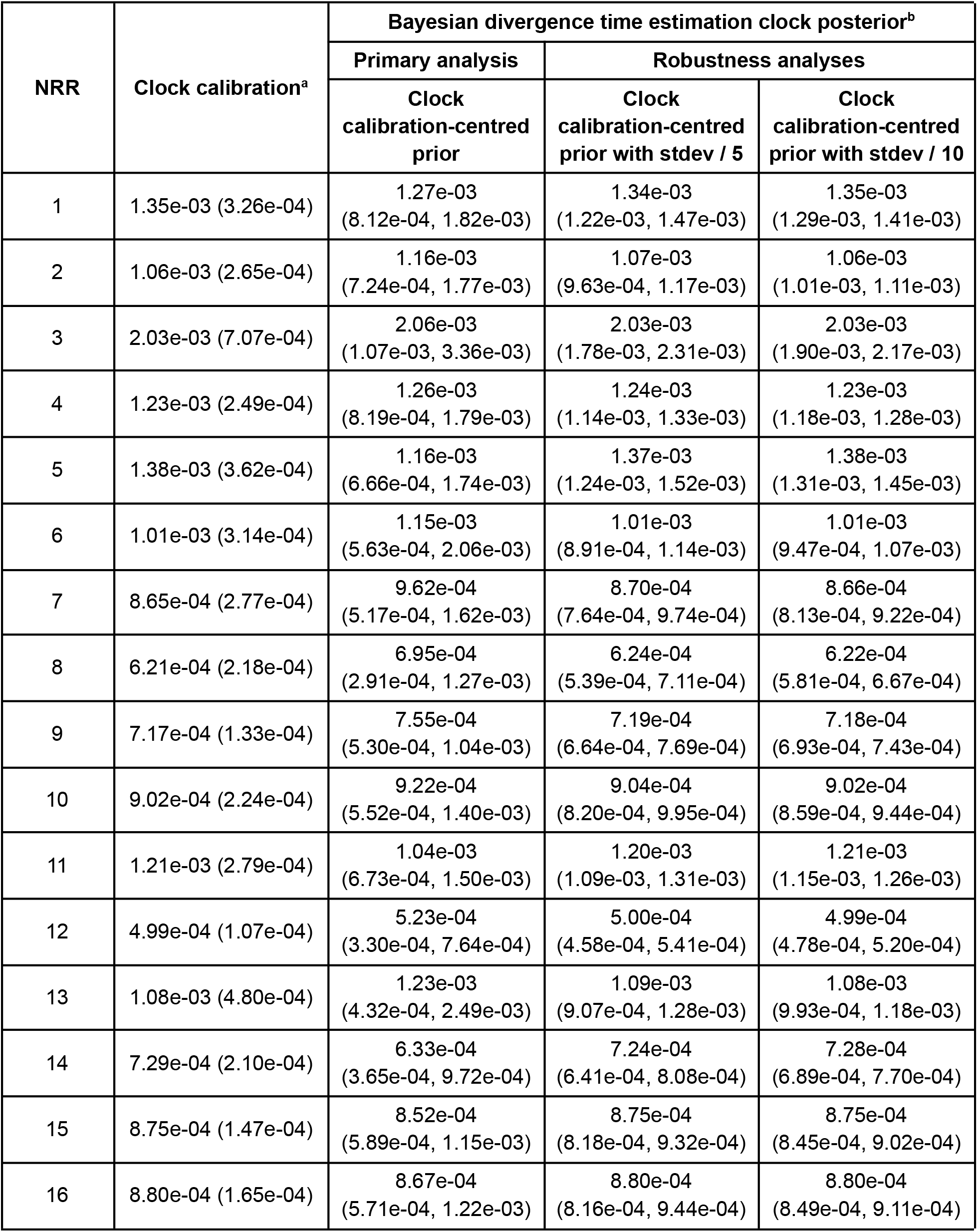
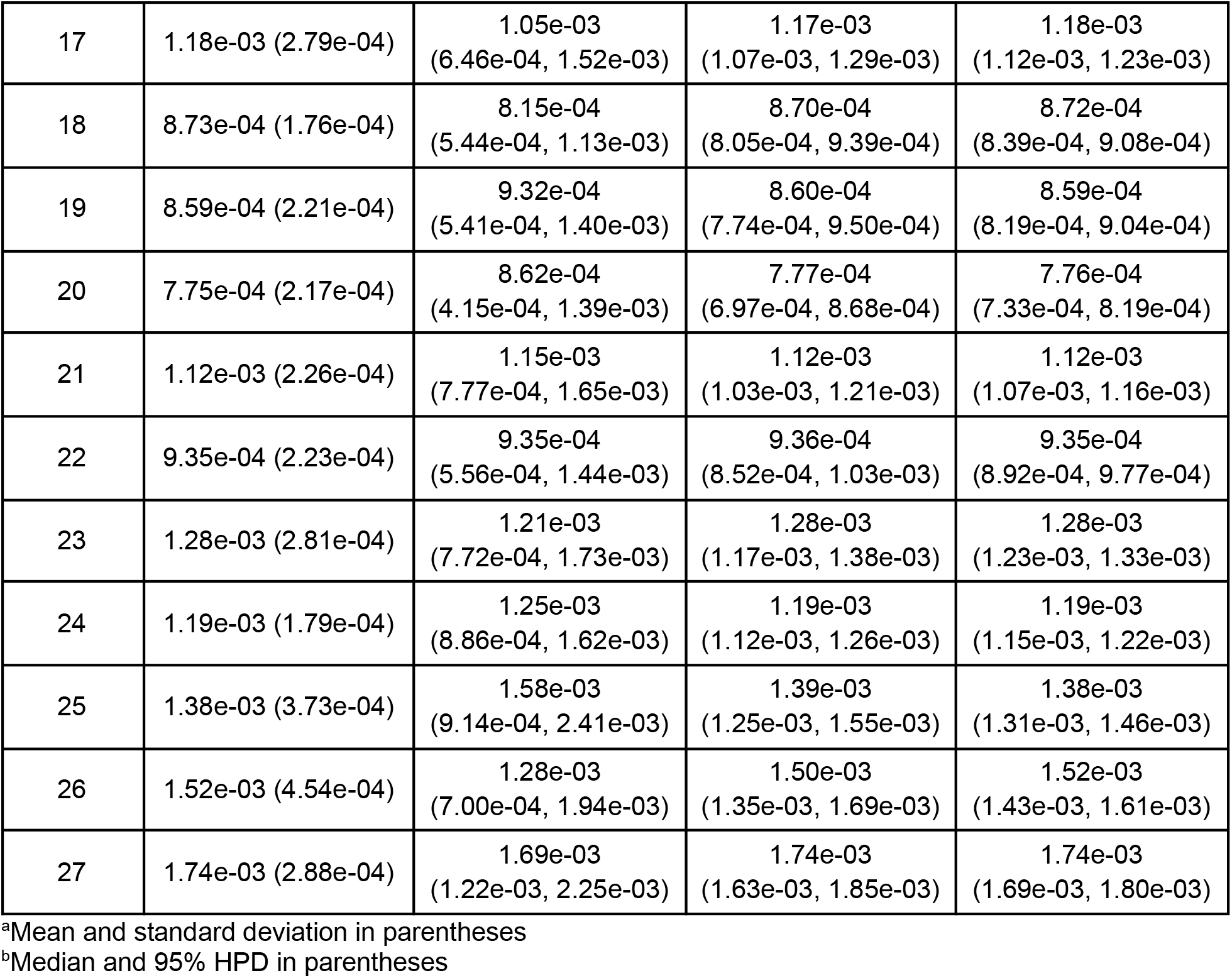
Posterior clock rates across each NRR from the clock calibration using SARS-CoV-2 genomes and Bayesian divergence time analyses of SARS-CoV-2-like viruses. The clock calibration posterior results are used as a prior for the Bayesian divergence time analyses, with the standard deviation of the clock calibration results varying across the analyses.

**Supplementary Table 3.** Posterior times (median and 95% HPD) of the closest-inferred ancestor of SARS-CoV-1 across each NRR. The SARS-CoV-1 clock calibration posterior serves as a prior for the analyses, with the standard deviation of the clock calibration results varying across the analyses and one analysis having an one additional civet SARS-CoV-1 genome and one additional human SARS-CoV-1 genome.

| NRR | Primary analysis | Robustness analyses |  |  |
| --- | --- | --- | --- | --- |
|  | Clock calibration-centred prior | Clock calibration-centred prior with stdev / 5 | Clock calibration-centred prior with stdev / 10 | Clock calibration-centred prior with additional genomes |
| 1 | 1996 (1990-2000) | 1998 (1994-2001) | 1998 (1994-2001) | 1995 (1989-2000) |
| 2 | 1992 (1980-2000) | 1996 (1991-2001) | 1997 (1991-2001) | 1991 (1978-1999) |
| 3 | 1991 (1972-2001) | 1991 (1974-2001) | 1991 (1975-2001) | 1987 (1969-1999) |
| 4 | 1993 (1982-2002) | 1998 (1991-2002) | 1998 (1991-2002) | 1993 (1981-2000) |
| 5 | 1994 (1987-2000) | 1997 (1990-2001) | 1997 (1991-2001) | 1994 (1986-2000) |
| 6 | 1999 (1993-2002) | 2000 (1997-2002) | 2001 (1997-2002) | 2001 (1997-2002) |
| 7 | 2001 (1997-2002) | 2001 (1998-2002) | 2001 (1998-2002) | 1999 (1994-2002) |
| 8 | 1985 (1960-2000) | 1999 (1994-2002) | 1999 (1994-2002) | 1984 (1960-1999) |
| 9 | 1984 (1962-2001) | 1987 (1960-2002) | 1988 (1971-2001) | 1976 (1952-1996) |
| 10 | 1999 (1993-2002) | 2000 (1994-2002) | 2000 (1994-2002) | 1997 (1990-2002) |
| 11 | 2000 (1995-2002) | 2001 (1998-2002) | 2001 (1998-2002) | 1999 (1994-2002) |
| 12 | 1995 (1986-2001) | 1999 (1994-2002) | 1999 (1994-2002) | 1997 (1989-2001) |
| 13 | 1997 (1986-2002) | 2000 (1995-2002) | 2000 (1995-2002) | 1997 (1987-2001) |
| 14 | 2001 (1998-2002) | 2001 (1999-2002) | 2001 (1999-2002) | 2000 (1995-2002) |
| 15 | 1992 (1977-2001) | 1969 (1930-1998) | 1974 (1930-2001) | 1970 (1925-1997) |
| 16 | 1999 (1993-2002) | 2000 (1997-2002) | 2000 (1995-2002) | 1998 (1991-2002) |
| 17 | 1997 (1991-2001) | 1998 (1994-2002) | 1992 (1980-2001) | 1991 (1977-2000) |
| 18 | 1989 (1971-2000) | 1986 (1961-1999) | 1988 (1969-1999) | 1989 (1972-1999) |
| 19 | 1963 (1934-1985) | 1980 (1963-1992) | 1980 (1964-1991) | 1963 (1934-1984) |
| 20 | 1978 (1953-1998) | 1985 (1966-2001) | 1986 (1965-2001) | 1979 (1952-1997) |
| 21 | 1977 (1948-1995) | 1989 (1967-2001) | 1989 (1968-2001) | 1975 (1947-1995) |
| 22 | 1979 (1940-2000) | 1988 (1963-2001) | 1989 (1962-2002) | 1980 (1952-1997) |
| 23 | 1991 (1966-2002) | 1995 (1975-2002) | 1994 (1975-2002) | 1989 (1941-2002) |
| 24 | 1999 (1989-2002) | 2000 (1992-2002) | 2000 (1992-2002) | 1996 (1969-2002) |
| 25 | 1999 (1995-2002) | 2000 (1996-2002) | 2000 (1996-2002) | 1998 (1994-2001) |
| 26 | 2000 (1995-2002) | 2001 (1997-2002) | 2001 (1997-2002) | 1996 (1986-2002) |
| 27 | 2000 (1994-2002) | 2000 (1996-2002) | 2000 (1997-2002) | 1998 (1989-2001) |
| 28 | 1999 (1992-2002) | 1999 (1994-2002) | 1999 (1994-2002) | 1998 (1990-2002) |
| 29 | 1996 (1986-2002) | 1998 (1990-2002) | 1998 (1991-2002) | 1994 (1984-2001) |
| 30 | 1989 (1945-2002) | 1996 (1971-2002) | 1996 (1969-2002) | 1982 (1863-2001) |
| 31 | 2001 (1999-2002) | 2001 (2000-2002) | 2001 (2000-2002) | 2000 (1997-2002) |

**Supplementary Table 4.** Posterior times (median and 95% HPD) of the closest-inferred ancestor of SARS-CoV-2 across each NRR. The SARS-CoV-2 clock calibration posterior serves as a prior for the analyses, with the standard deviation of the clock calibration results varying across the analyses.

| NRR | Primary analysis | Robustness analyses |  |
| --- | --- | --- | --- |
|  | Clock calibration-centred prior | Clock calibration-centred prior with stdev / 5 | Clock calibration-centred prior with stdev / 10 |
| 1 | 2011 (2006-2015) | 2012 (2007-2015) | 2012 (2008-2015) |
| 2 | 2013 (2007-2017) | 2012 (2007-2016) | 2012 (2008-2016) |
| 3 | 2017 (2014-2019) | 2017 (2014-2019) | 2017 (2014-2019) |
| 4 | 2010 (2000-2017) | 2010 (2002-2016) | 2010 (2002-2017) |
| 5 | 2013 (2008-2017) | 2014 (2010-2018) | 2014 (2010-2017) |
| 6 | 2015 (2009-2019) | 2015 (2009-2018) | 2015 (2010-2019) |
| 7 | 2014 (2008-2018) | 2013 (2009-2017) | 2013 (2009-2017) |
| 8 | 2004 (1979-2016) | 2002 (1984-2014) | 2003 (1984-2014) |
| 9 | 1994 (1978-2004) | 1992 (1981-2001) | 1992 (1981-2001) |
| 10 | 2008 (1997-2016) | 2008 (1999-2016) | 2008 (1997-2015) |
| 11 | 2010 (2004-2014) | 2011 (2007-2014) | 2011 (2007-2014) |
| 12 | 1997 (1976-2011) | 1996 (1979-2009) | 1996 (1979-2009) |
| 13 | 2010 (1984-2019) | 2010 (1989-2019) | 2010 (1991-2019) |
| 14 | 1997 (1982-2007) | 1999 (1991-2005) | 1999 (1991-2005) |
| 15 | 2012 (2007-2015) | 2012 (2008-2015) | 2012 (2008-2015) |
| 16 | 2003 (1993-2010) | 2003 (1996-2009) | 2003 (1995-2008) |
| 17 | 2012 (2006-2017) | 2013 (2008-2017) | 2013 (2008-2017) |
| 18 | 2004 (1996-2010) | 2005 (2000-2009) | 2005 (1999-2009) |
| 19 | 2012 (2005-2017) | 2011 (2006-2016) | 2011 (2006-2016) |
| 20 | 2004 (1988-2014) | 2002 (1991-2011) | 2002 (1991-2011) |
| 21 | 1992 (1977-2003) | 1991 (1980-2000) | 1991 (1980-1999) |
| 22 | 1982 (1946-2007) | 1982 (1955-2004) | 1982 (1955-2004) |
| 23 | 1989 (1969-2007) | 1991 (1978-2006) | 1990 (1977-2005) |
| 24 | 1992 (1981-2001) | 1991 (1984-1997) | 1991 (1983-1997) |
| 25 | 2006 (1993-2015) | 2004 (1993-2013) | 2004 (1994-2013) |
| 26 | 2009 (2000-2016) | 2010 (2004-2016) | 2010 (2004-2016) |
| 27 | 2013 (2008-2016) | 2013 (2008-2016) | 2013 (2008-2016) |

**Supplementary Table 5.** Comparison of the weighted diffusion coefficient and isolation-by-distance signal estimated for different data sets of viral genomes. The isolation-by-distance (IBD) signal has been estimated by computing the Spearman correlation coefficient (rS) between the patristic and great-circle geographic distances computed for each pair of virus samples. For each data set and metric, we report both the posterior median estimate and the 95% HPD interval. (*) estimates based on the combined analysis of lineages L1 and L3. (**) estimates obtained while only considering phylogenetic branches occurring less than 50 years ago and bat samples, but similar results are obtained when considering branches occurring less than 75 years ago (2839 km^2^/year [1595, 4558] for SARS-CoV-1-like viruses, and 2536 km^2^/year [638, 6089] for SARS-CoV-2-like viruses) or when considering all samples for SARS-CoV-1-like viruses (3030 km^2^/year [1965, 4668]). When considering all samples for SARS-CoV-2-like viruses, we however obtain a higher estimate for the weighted diffusion coefficient, but with heavily overlapping confidence intervals and the same order of magnitude (3817 km^2^/year [1824, 6929]). SARS-CoV-2-like viruses (bats)

| Data set | Weighted diffusion coefficient (km<sup>2</sup>/year) | IBD signal ($r_S$) | Number of samples | Reference |
| --- | --- | --- | --- | --- |
| Porcine deltacoronavirus, China | 26093 [17312, 37020] | -0.02 [-0.07, 0.05] | 97 | He et al. (2020) |
| Getah virus, China | 24930 [17612, 35504] | 0.06 [0.00, 0.14] | 125 | Zhao et al. (2023) |
| West Nile virus, North America | 58757 [55115, 62060] | 0.00 [-0.02, 0.03] | 801 | Dellicour et al. (2020) |
| Lassa virus, segment S, Africa | 42 [38, 49] | 0.68 [0.64, 0.72] | 254 | Klitting et al. (2022) |
| Rabies virus (bats)*, Peru | 1416 [1105, 1852] | 0.73 [0.55, 0.83] | 260 | Streicker et al. (2016) |
| Rabies virus (skunks), USA | 580 [475, 676] | 0.49 [0.48, 0.60] | 241 | Kuzmina et al. (2013) |
| SARS-CoV-1-like viruses (bats) | 2884 [1654, 4358]** | 0.45 [0.36, 0.59] | 134 | (present study) |
| SARS-CoV-2-like viruses (bats) | 2050 [233, 5470]** | 0.66 [0.12, 0.83] | 20 | (present study) |

